# Toward Unified Biomarkers for Focal Epilepsy

**DOI:** 10.1101/2025.04.15.647522

**Authors:** Sheng H Wang, Paul Ferrari, Gabriele Arnulfo, Morgane Marzulli, David Degras, Vladislav Myrov, Satu Palva, Lino Nobili, Philippe Ciuciu, J Matias Palva

**Author notes:** Correspondence to: Sheng H. Wang, Full address: CEA, Joliot, NeuroSpin, Bât. 145, Point Courrier 156, 91191, Gif-sur-Yvette cedex, France. These authors contributed equally to this work (Philippe Ciuciu, J Matias Palva). All authors declare no competing interests. The funders had no role in study design, data acquisition, analyses, decision to publish, and preparation of the manuscript.

## Abstract

Accurately localizing the epileptogenic network (EpiNet) remains a major barrier to effective epilepsy treatment, largely due to limited mechanistic understanding. The EpiNet is a patient-specific brain network shaped by complex, overlapping pathology. While combining biomarkers can improve localization, it also generates high-dimensional feature data that increases the risk of overfitting and reduces interpretability. We hypothesized that the core epileptogenic dynamics could be captured in a low-dimensional latent space derived from empirical data, without the need to record seizures. From interictal stereo-EEG (SEEG) recordings in 64 patients (29 females), we extracted 260 neuronal features and reduced them to 10 latent components using singular value decomposition. A classifier trained on these 10 components was then simplified into a probabilistic EpiNet model requiring only two components as input. Individual position in this two-dimensional latent space correlated with previously reported classification accuracy (*r*^2^=0.5), supporting its functional relevance. In three independent patients, the probabilistic model captured time-varying epileptogenic dynamics during sleep-SEEG recordings, corroborated clinical assessments, and achieved peak classification accuracies of 0.63, 0.85, and 0.94. These predictions were independently validated by tensor component analysis. Together, these results provide evidence for a robust low-dimensional representation of epileptogenicity across brain states and pathological substrates. This approach simplifies interpretation, facilitates integration of additional biomarkers, and enables large-scale cohort analyses, establishing a proof of concept for a unified framework for epilepsy biomarkers.

**Significance Statement:** To advance mechanistic understanding of large-scale brain dynamics underlying epilepsy, we combined novel epilepsy biomarkers with interpretable machine learning. From interictal SEEG, we extracted 260 connectivity and criticality features. Dimensionality reduction of these raw features showed that only two components were needed to identify epileptogenic networks, reducing the feature space by >99%. These components were highlighted by abnormal power-law scaling, bistability, and strong inhibition or excitation in the 15–200 Hz range, consistent with prior findings. The model tracked neuropathological dynamics over hours and was validated through tensor component analysis, suggesting that epileptogenic activity is dynamic, subject-specific, and sparsely represented in both state space and cortical networks.

## Introduction

Epilepsy affects 1% of the general population, and one in three patients are drug-resistant (DRE)^1^. The most effective treatment for gaining seizure control in focal onset DRE is surgical resection or disconnection of the seizure-causing brain tissue known as the epileptogenic zone (EZ). A growing body of recent research^2–5^ suggests that, rather than a single discrete EZ or seizure focus, multiple overlapping neuropathological substrates form an epileptogenic network (EpiNet) that are collectively responsible for generating seizures. The EpiNet may comprise structural lesion^6^, high frequency oscillations zone^7^, irritative (spiking) zone^8^, and seizure zone, where seizures are clinically observed^3^. Various biomarkers have been proposed to localize these epileptogenic mechanisms^9,10^. However, surgical outcomes vary significantly, with 30% to 70% patients experiencing seizure recurrence within few months of surgery^11,12^, suggesting that accurate EpiNet localization remains a current challenge.

We recently advanced novel epilepsy biomarkers inspired by complex systems theory and the brain criticality hypothesis^13–15^. We proposed that the brain functions optimally when operating in a critical regime between order and disorder^14,16^. Empirical evidence linking aberrant brain criticality to epilepsy includes abnormal power-law scaling^13,17^, strong inhibition^18,19^, and high bistability in neuronal oscillations^18,20^ that suggests a shift towards catastrophic events like that in many complex systems^15,21^. Aberrant local criticality is believed to coexist with a large-scale trend towards supercriticality, characterized by elevated phase synchrony^14^ and cross-frequency phase-amplitude coupling^22^ within and around the EpiNet. Utilizing both criticality and large-scale synchrony biomarkers has yielded more accurate localization compared to individual biomarkers^18^, offering support for the multi-component hypothesis for the EZ^3^.

However, employing multiple biomarkers to assess dozens of narrow-band oscillations leads to high dimensional feature sets, posing a major challenge to advancing automated, data driven localization. Having hundreds or thousands of features requires enormous computational resources and makes results interpretation difficult between physicians. It also risks over-fitting and prevents efficient training of machine learning models. This impedes the addition of novel^23,24^ biomarkers or established clinical biomarkers, such as high-frequency oscillations, slow waves, and spikes, thus hindering the efforts to study their functional relationships and to understand why clinical biomarkers often lose specificity in practice^25^. Moreover, the classifiers used earlier have high overhead and may handle thousands of sampled brain regions but would become impractical for studying larger clinical data. Therefore, we aimed to advance a new framework that would simplify feature interpretation and resolve aforementioned technical challenges.

We propose that the EpiNet can be identified using a “latent space” composed of only a few principal components derived from dimensionality reduction of cohort-level interictal neuronal features. We previously showed that the high-dimensional feature sets decomposed in this manner could identify individual EpiNet^26^. Here, we speculate that this computationally tractable framework enables a time-resolved assessment of an individual’s EpiNet state, such that its expression, or “saliency”, in the latent space fluctuates over time as recently proposed ^27–29^ (Fig. 1B–C). We further speculate that temporal saliency correlates with increased seizure propensity, or “Seizure Risk,” as reflected in epileptiform spikes, and that this saliency should be positively correlated with individual classification accuracy based on raw features^18^. To test this Saliency Model, we applied the framework from our previous study^26^ to 30 hours of sleep SEEG recordings from three independent epilepsy patients.

**Figure 1.**
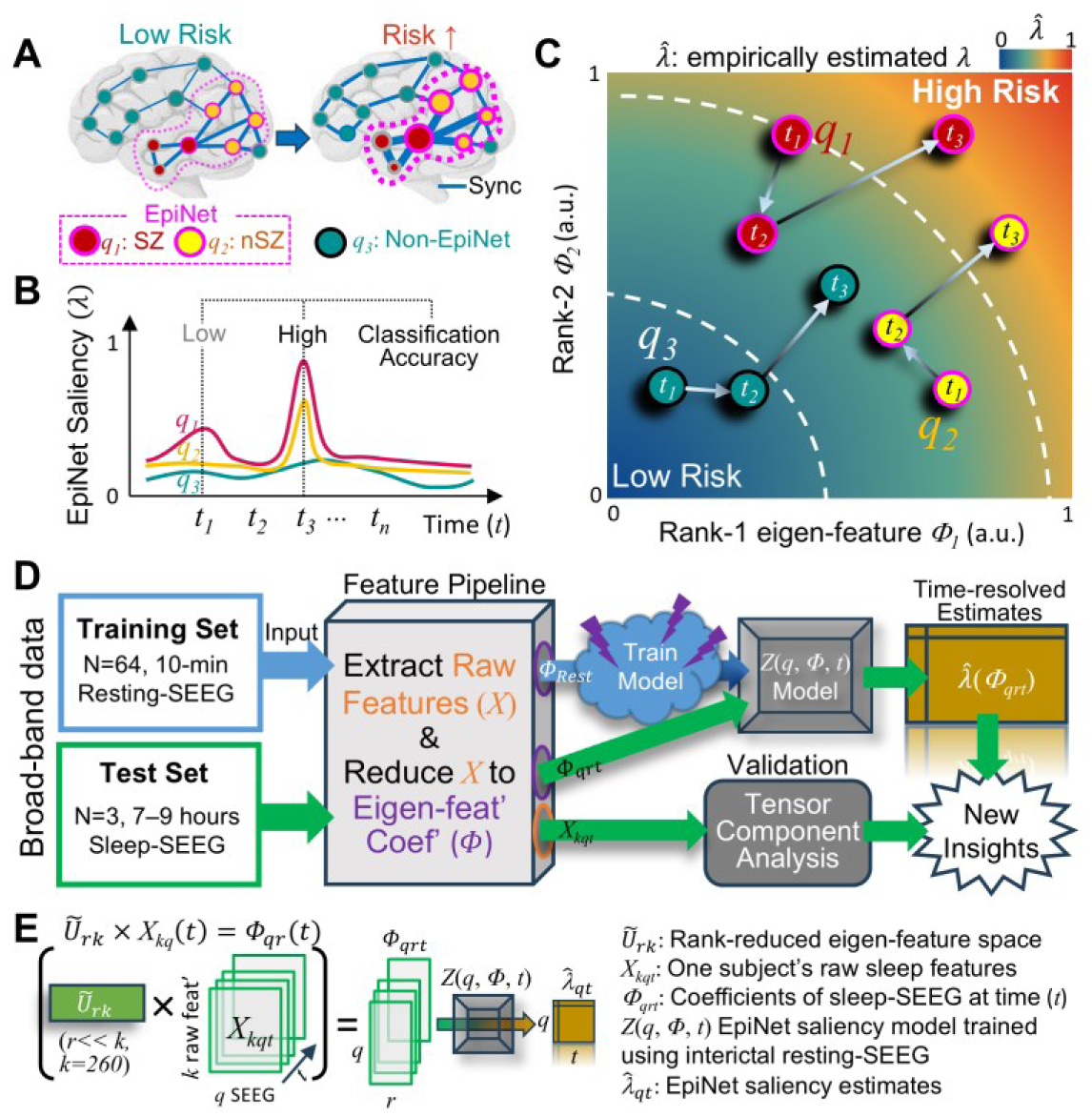
Study design. **(A–C)** The hypothesis. **(A)** An increased seizure risk is associated with aberrant local (nodes) criticality and strong synchrony (edges) within and around the epileptogenic network (EpiNet). **(B)** As seizure risk increases, the EpiNet, including its constituent seizure zone (SZ) and non-SZ (nEZ), exhibits more salient pathological features (larger *λ*) than other regions (*q_3_*). The saliency of the EpiNet fluctuates over time (*t*). **(C)** High-dimensional interictal raw brain dynamics features of a training dataset are projected into a low-dimensional eigenfeature space, allowing classifiers to be trained to assess individual regions’ EpiNet-saliency (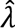). **(D)** The analysis pipeline. **(E)** Test the low-dimensional EpiNet-saliency model with sleep-SEEG. Matrix multiplication (×): project one subject’s raw sleep features *X*_*kq*_(*t*) into the eigenfeature space *Ũ*_*rk*_derived from interictal-SEEG. The resulting coefficients *Φ*_*qrt*_is used as input for the model. *q*: number of SEEG contacts; *r*: reduced rank<< raw feature number *k* = 260; *t*: number of observation windows with 10-minute resolution.

## Materials and methods

### Overview

Two hundred sixty SEEG features of the training set were reduced to ten eigenfeatures under the hypothesis that this reduced dimensional feature set, or ‘Latent Space’, is capable of differentiating the seizure zone (SZ) from where seizures activity was identified by physicians. These eigenfeatures were used to train a classifier that identified EpiNet contacts—those with epileptogenic characteristics regardless of clinical SZ or non-SZ labels. From this, we derived the EpiNet-saliency model, a simpler probabilistic framework using only two eigenfeatures to estimate EpiNet likelihood. To test generalizability, sleep-SEEG from an independent cohort was linearly projected into the interictal-SEEG eigenfeature space and evaluated with the saliency model (Fig. 1E). Model assessments were compared against clinically identified cortical dysplasia and further validated with tensor component analysis.

### Subjects and recording

The training set included 7,183 SEEG contacts from 64 patients (mean ± SD age: 29.7 ± 9.5, 29 females) with focal onset seizures treated at the Niguarda “Ca’ Granda” Hospital, Milan, Italy. They underwent SEEG for the first time and had no prior epilepsy surgery. Ten-minute of eyes-closed, interictal brain activity were recorded with a 192-channel SEEG amplifier system (NIHON-KOHDEN NEUROFAX-1100) at a sampling rate of 1 kHz, ensuring no seizure activity at least one hour before and after these recordings^30^. The test set included three patients from a separate cohort treated at the same institute. Their epilepsy was caused by type-2 focal cortical dysplasia (FCD), confirmed by post-surgical histopathological study. Structural MRI and post-implant CT data were unavailable, and a bipolar referencing scheme was used for them. All training and test subjects were under anti-seizure medication.

The time from the last drug administration to SEEG recording was not controlled, and the drug effects on individual epileptogenic features therefore were not considered. All subjects gave written informed consent for participation in research studies and for publication of results pertaining to their data. This study was approved by the ethical committee (ID 939) of the Niguarda “Ca’ Granda” Hospital and was performed according to the Declaration of Helsinki.

### Clinically identified seizure zone (SZ)

Physicians identified SZ by visual inspection of SEEG traces. In focal epilepsy patients, seizures often originate from a well-defined seizure onset zone (SOZ), characterized by abnormal electrophysiological activity such as rapid low-voltage bursts, spike-wave discharges, and slow bursts with poly-spikes. Other regions within the pathological brain network, known as the seizure propagation zone (SPZ), do not initiate seizures but exhibit rhythmic changes shortly after seizure onset in the SOZ. In some cases, a brain region may be classified as both SOZ and SPZ, referred to as the seizure onset and propagation zone (SOPZ). For this study, we grouped SOZ, SPZ, and SOPZ under the broader term SZ. Any brain regions not identified as SZ were considered tentatively non-seizure zones (nSZ).

### Preprocessing and filtering

SEEG time series were low-pass filtered with FIR filter with cutoff at 440 Hz and stop-band at 500 Hz (60 Hz transition band, −6 dB suppression at 475 dB, maximal ripples in pass-band 2%). Fifty hertz line-noise and its harmonics were excluded with a band-stop FIR filter with 53 dB suppression and 1 Hz band-stop widths. We excluded SEEG contacts (on average 1.3 ± 1.2 with range 0–10) that showed non-physiological activity. On average these subjects had 113±16.2 (±SD) contacts recorded from grey-matter, and the recordings were referenced using the nearest white-matter contact, a referencing scheme that yields signals with a consistent polarity, attenuates signal mixing, and provides more accurate phase estimates^31^. Criticality features in local oscillations were assessed for 20 narrow-band frequencies obtained with Morlet wavelets (*m*=5), ranging from 2 to 225 Hz with equal log_10_ spacing^15,18^. All-to-all phase synchrony was assessed across 50 narrow-band frequencies with Morlet wavelets (*m*=7.5), ranging from 2 to 450 Hz with equal log₁₀ spacing^14,30^.

### Criticality and synchrony features

Features were assessed for narrow-band oscillations following previous work^14,15,18^, with formal definitions provided in the Supplementary Methods. Local criticality features include detrended fluctuation analysis (DFA) exponent, bistability index (BiS), and functional E/I balance (fE/I). Large-scale synchrony was assessed using the phase-locking value (PLV), from which adjacency matrices were derived to compute synchrony features.

DFA quantifies long-range temporal correlations (LRTCs) in a time series and peaks inside an extended critical-like regime^32^ known as the Griffiths phase^14^ (Fig. 2A). Outside this regime, LRTCs approach white noise level. Significant LRTCs (DFA > 0.6) are commonly used to indicate operation within the Griffiths phase. Within this regime, the fE/I index characterizes three functional profiles—Inhibition-dominant, balanced, and Excitation-dominant—based on correlation between time-resolved amplitudes and DFA^33,34^(Fig. 2A).

**Figure 2.**
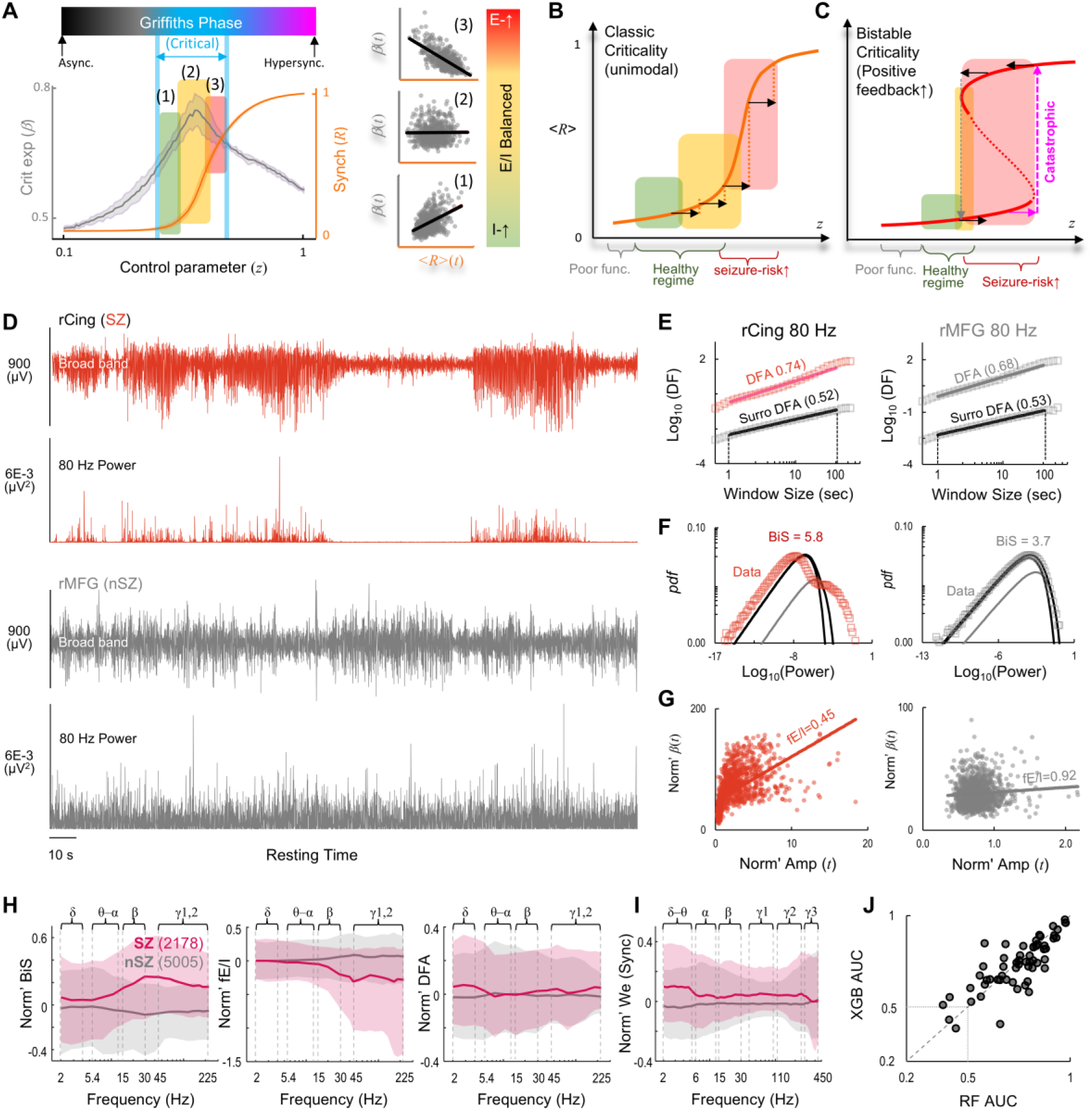
Feature overlap between seizure zone (SZ) and non-SZ (nSZ) introduces variability in accuracy of supervised SZ-classification. (A–C) Theoretical motivation for the biomarkers (Methods). (A) Three excitation/inhibition (E/I) profiles inside an extended critical regime (Griffiths phase) based on time-resolved analysis. In insets, Markers indicate temporal samples. The neuronal dynamical regime of (B) classic criticality and (C) bistable criticality. (D) Broad-band and narrow-band time series of interictal-SEEG from a SZ and a nearby nSZ contact. rCing: right cingulate; rMFG: right middle frontal gyrus. (E–G) The fitting of criticality metrics for the time series from (D). (E) Computation of critical exponents of long-range temporal correlations (DFA), (F) bistability index (BiS), and (G) functional E/I estimate (fE/I). (H–I) Within-subject normalized (H) criticality and (I) synchrony features. We: effective weight of phase synchrony. Shaded areas: 25% and 75%-tile. (J) Supervised SZ-classification accuracy. AUC: area under receiver operating characteristics curve. Markers: subjects; RF (Random Forest) and XGB (XGBoost) are two supervised classifiers trained using clinically identified seizure zone contacts.

Neuronal oscillations may exhibit both unimodal or bistable distribution (Fig. 2B–C, respectively), likely caused by an underlying positive feedback^15,35^. When interictal recording durations are identical, high BiS index serves as strong predictors of catastrophic transitions toward supercritical, *e.g.* seizure-like dynamics^15,18^.

Synchrony features were extracted from narrow-band *PLV* adjacency matrices^18^. First-order features, *e.g.*, eigenvector centrality (EVC) and effective weight (We), characterize strongly connected hub regions. Second-order features, *e.g.*, clustering coefficient (Cc) and local efficiency (LE), identify regions whose neighbors are densely interconnected.

Criticality and synchrony features were assessed using an in-house Python toolbox and the <u>Brain Connectivity Toolbox</u>. For the training set subjects, features were extracted with a 10-minute interictal-SEEG recording (Fig. 1D). For the test set subjects, features were extracted from 7–9 hours of sleep-SEEG by applying a 10-minute sliding window with a one-minute step size. The time-resolved raw sleep features for each test subject form a three-way tensor (*X* ∈ *R*^*kxqxt*^), representing *k* features, *q* contacts, and *t* time points (Fig. 1E). The parameters for filtering and feature extraction were kept identical for both datasets.

### Supervised classifiers: replication of previous work

Based on observed spatial similarity in our previous studies^15,18^, twenty normalized narrow-band criticality estimates were averaged into four frequency clusters as δ (2–4 Hz), θ − α (5.4– 11 Hz), β (15–30 Hz), and γ1,2 (45–225 Hz). Similarly, fifty normalized narrow-band synchrony features were collapsed into six frequency clusters as: δ − θ (2–5.4Hz), α (6.1–13 Hz), β (15–30 Hz), γ1 (40–96 Hz), γ2 (110–250 Hz), and γ3 (270–450Hz). To optimally separate SZ and nSZ contacts within subjects, normalization was done for each subject as: *x*_*i*_ = (*x*_*i*_ − median(*X*))./max(*X* − median(*X*)), where *x*_*i*_represent a contact, *X* is a 1D vector of real numbers representing one narrow-band feature for all contacts. This computation resulted in 36 band-collapsed features to characterize each contact. For within-patient supervised SZ classification, we used leave-one-out validation (Fig. 2I), *i.e.*, a patient’s contacts served as the test set and the remaining subjects formed the training set. We previously used the Random Forest algorithm^36^. To ensure robustness, we employed an additional classifier eXtreme Gradient Boosting (XGBoost)^37^ from the scikit-learn toolbox^38^. Random Forest, previously used in our studies^15,18^, is a machine learning method that leverages bootstrapped training data and combines decision tree simplicity with flexibility in handling new data. XGBoost, based on gradient boosting, sequentially refines decision trees by correcting previous errors, offering high speed, accuracy, and built-in regularization to prevent overfitting.

### Spike density

Interictal epileptiform discharges (IEDs) are characterized by fast, high-amplitude transients lasting less than 200 ms^39^, distinguishable from the background, and often spreading across multiple brain regions^30^. The IEDs were identified following the approach described in^18^. Briefly, for a 10-minute recording session of interictal or sleep SEEG, we divided the broad-band signal from each SEEG contact into non-overlapping 0.5-second windows. A window was classified as ‘spiky’ if at least three consecutive samples exceeded seven times the standard deviation above the contact’s mean amplitude. The spike density of a contact was then defined as the percentage of windows classified as spiky over the 10-minute session.

### The EpiNet-saliency model

#### Model design

An increase in seizure risk from baseline should be associated with multiple raw feature patterns such as strong synchrony and aberrant local criticality, most pronounced within and around the EpiNet as previously reported^15,18,26^ (Fig. 1A). If the low-dimensional epileptogenicity hypothesis holds, a brain region’s pathological feature saliency over a time period could be characterized by its trajectory in a low-dimensional state space, *i.e.*, a graph that shows all the possible ways a neuronal system may behave (Fig. 1B–C). The EpiNet-saliency model was designed to be a lookup table using the two most predictive eigenfeatures as inputs, eliminating the need for retraining with new data and improving efficiency. Moreover, as seizure risk is known to fluctuate over hours^26–29^, prolonged recordings are desirable to capture a full spectrum of the EpiNet-saliency.

#### Step 1: obtain an eigenfeature space

Singular value decomposition (SVD) of the raw interictal features is defined as:

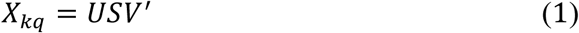

where raw features *X*_*kq*_ ∈ *R*^*k×q*^ is a 2D array with each element represents *k*-th feature of *q*-th contact (*k* = 260 features, *q* = 7,183 contacts from 64 subjects); *U* ∈ *R*^*k*×*k*^is a unitary matrix representing the full-rank eigenfeature space, with each of its column representing linear combination of raw-features (an example see Fig. 3C and Supplementary Fig. 1); *S* ∈ *R*^*k×q*^ is a rectangular diagonal matrix with singular values, and *V*^′^ ∈ *R*^*q*×*q*^ is a unitary matrix. Neither *S* nor *V*′ are used.

**Figure 3.**
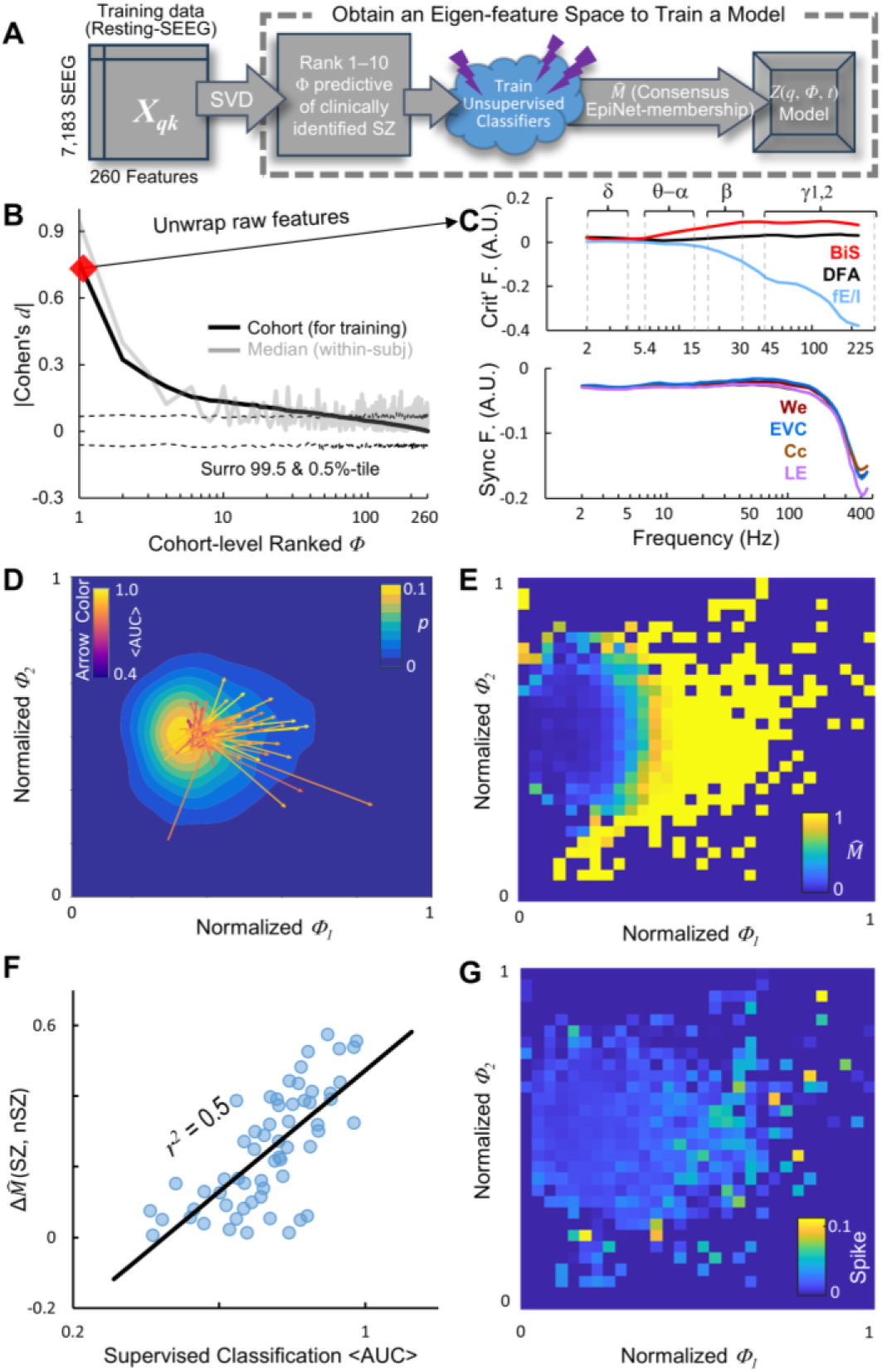
Obtain eigenfeature coefficients to train a model to assess EpiNet-saliency. **(A)** Pipeline. **(B)** Eigen feature coefficient ranked by cohort-level effect size of difference between SZ and nSZ contacts (black). Validation was performed using the median of within-subject effect size of differences (gray), *i.e.*, a point on the median line indicates the smallest effect size observed for the corresponding coefficient in at least half of the subjects. Dashed lines: confidence intervals observed from 10^4^ SZ-nSZ label shuffled surrogates. **(C)** Normalized features of the rank-1 eigenfeature. **(D)** Cohort level joint distribution. Arrows: subjects; arrowhead: SZ; arrow-tail: nSZ. **(E)** Joint-distribution of EpiNet-membership (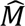). For each a pair of (*Φ_1_*, *Φ_2_*), multiple contacts may exist; their mean 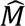 is shown as the pixel color. **(F)** Correlation between supervised SZ-classification accuracy (from Fig 2I) and the within-subject difference in 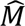 between SZ and nSZ. **(G)** Joint distribution of mean contact spikes.

#### Step 2: seizure-zone relevant eigenfeature selection

The raw feature *X*_*kq*_ of the training data is projected into the full-rank eigenfeature space *U*:

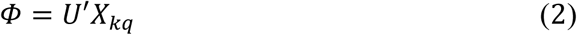

where the eigenfeature coefficients *Φ* ∈ *R*^*k×q*^ represents the projection of raw features onto *U*. For each coefficient, we assessed the effect size of the difference between SZ and nSZ contacts at the cohort level using Cohen’s *d*. Coefficients were then ranked by Cohen’s *d* (Fig. 3B), with larger *d* values indicating greater separation between SZ and nSZ contacts. This cohort-level result was validated by within-subject effect size analysis (gray line, Fig. 3B). To construct a reduced representation, we retained the ten coefficients with the largest effect sizes relative to surrogate data, yielding a rank-reduced *Ũ* ∈ *R*^*r*×*k*^with *r* =10. This defines an epileptogenicity-relevant sub-space of *U* that preserves 10 of 260 of its original dimensions.

#### Step 3: train classifiers to identify the EpiNet

We speculate that the EpiNet includes both clinically identified SZ and pathological nSZ contacts (Fig. 1A), and we aimed to distinguish EpiNet from non-EpiNet. Given the individual variability and potential functional overlap among brain regions^3^, we selected soft-partitioning classifiers—Gaussian mixture models (GMM)^38^ and Fuzzy C-means (FCM)^40^—over hard-partitioning classifiers used in previous studies^15,18^. These two classifiers, despite their reliance on different assumptions, should yield correlated accuracy if the latent features reliably identify EpiNet samples. We classified contacts pooled across training subjects. To examine how the number of components influenced membership estimates, we tested both classifiers incrementally using 2 to 10 components ranked by eigenfeature coefficients *Φ* (Fig. 3B). In each test, the classifiers assigned each contact an EpiNet membership (*M* ∈ {0, 1}), where a value close to 1 indicating high likelihood of belonging to the EpiNet. Across the tests, *M* estimates were consistent (Supplementary Fig. 2A) and were therefore averaged within each classifier. These mean *M* from GMM and FCM were strongly correlated (Pearson’s *r* = 0.893, -log_10_(*p*) > 12, Supplementary Fig. 4B), supporting the validity of using a *consensus EpiNet membership* (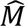) by averaging the two means. Contacts with 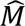 near 1 is highly likely to belong to the EpiNet, while those near 0 are unlikely.

#### Step 4: fitting the EpiNet-saliency model

We next computed joint distribution for <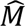> in rank-1 and rank-2 EpiNet-relevant coefficient (*Φ_1_–Φ_2_*) space, computed using 30 linear bins for both *Φ_1_* and *Φ_2_* (Fig. 3E). To make it a complete lookup table for all possible (*Φ_1_–Φ_2_*) combinations in new data, we fitted it as a smooth surface using MATLAB’s <u>Curve Fitting Toolbox</u> with ‘linear interpolant’ method to approximate values between known data points and with ‘linear extrapolation’ for other missing data points. The resulting fitted surface represents the EpiNet-saliency assessment 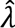 ∈ [0, 1] for a given contact in the (*Φ_1_–Φ_2_*) coordinate space, where 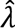 =1 indicates maximal saliency (Fig. 4A). To assess the assessment for a new contact, MATLAB function fitresult() is used by providing its *Φ_1_* and *Φ_2_* as inputs.

**Figure 4.**
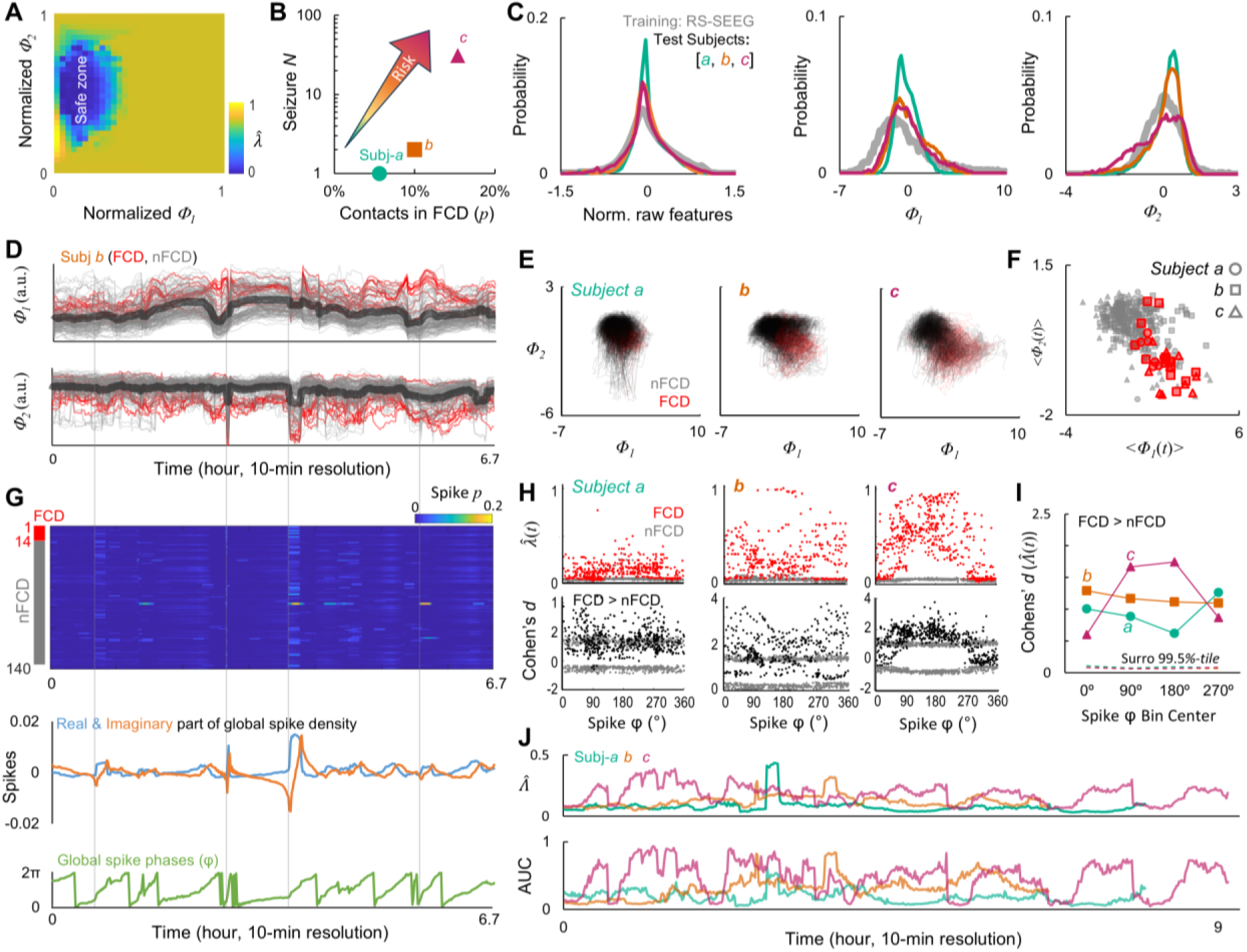
Cross-domain validate the model. **(A)** The EpiNet-saliency model visualized as a 2-dimensional lookup table. **(B)** Total number of seizure events observed during pre-surgical monitoring as a function of contacts inside dysplastic cortices (FCD). **(C–F)** Properties of *Φ_1_* and *Φ_2_*. **(C)** Probability distribution of normalized raw features, *Φ_1_*, and *Φ_2_* of the training and test subjects, with all features of all contacts pooled. **(D)** One subject’s individual contacts and mean (thick) *Φ_1_* and *Φ_2_* over time with temporal resolution of 10-minute. **(E)** All contacts’ trajectory overlaid. **(F)** Time-averaged *Φ_1_* and *Φ_2_*; subjects’ contacts are identified by marker shape defined in **(B)**. **(G)** Spikes observed from the same subject’s recording whose (*Φ_1_*, *Φ_2_*) are shown in (D). Top: contact spike density; middle: real (Re) and imaginary (Im) part of mean spike density across contacts; bottom: phases of mean spike density. **(H)** Top: distributions of EpiNet-saliency as a function of spike phases; bottom: distributions of effect sizes of differences between FCD and nFCD in EpiNet-saliency as a function of spike phases, gray markers represent surrogate observation. **(I)** Effect sizes in four spike phase bins. Dashed line: confidence interval observed from 10^3^ label-shuffled surrogates. **(J)** Top: time-resolved global EpiNet-saliency (mean across contacts); bottom: area under receiver-operating characteristic curve.

### Test the EpiNet-saliency

For each test subject, time-resolved eigenfeature coefficient *Φ*_*qrt*_was obtained by projecting (Eq. 2) the raw sleep features *X*_*kqt*_into the rank-reduced eigenfeature space *Ũ*_*rk*_of the interictal-SEEG (Fig. 1E). The rank-1 and rank-2 EpiNet-relevant coefficients from *Φ*_*qrt*_was then used as the input to the saliency model that returns an estimate 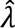(*q*, [*Φ*_1_, *Φ*_2_], *t*) for contact *q* observed at time *t.* Thus, given rank-1 and −2 coefficients [*Φ_1_*, *Φ_2_*] of a new contact *q* measured for time window *t*, the model returns an assessment 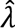(*q*, *t*). This can be done each contact across all the time windows. The average 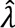 across contacts at time *t* characterizes a momentary global EpiNet-saliency 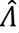(*t*) for time *t*.

### Tensor component analyses

The canonical polyadic tensor decomposition^41^ breaks down raw sleep features (*X*_*kqt*_), a three-way tensor, into a sum of rank-1 tensors, making it easier to analyze the hidden patterns in each of the three modes (Fig 5A left):

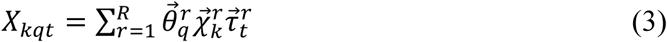

**Figure 5.**
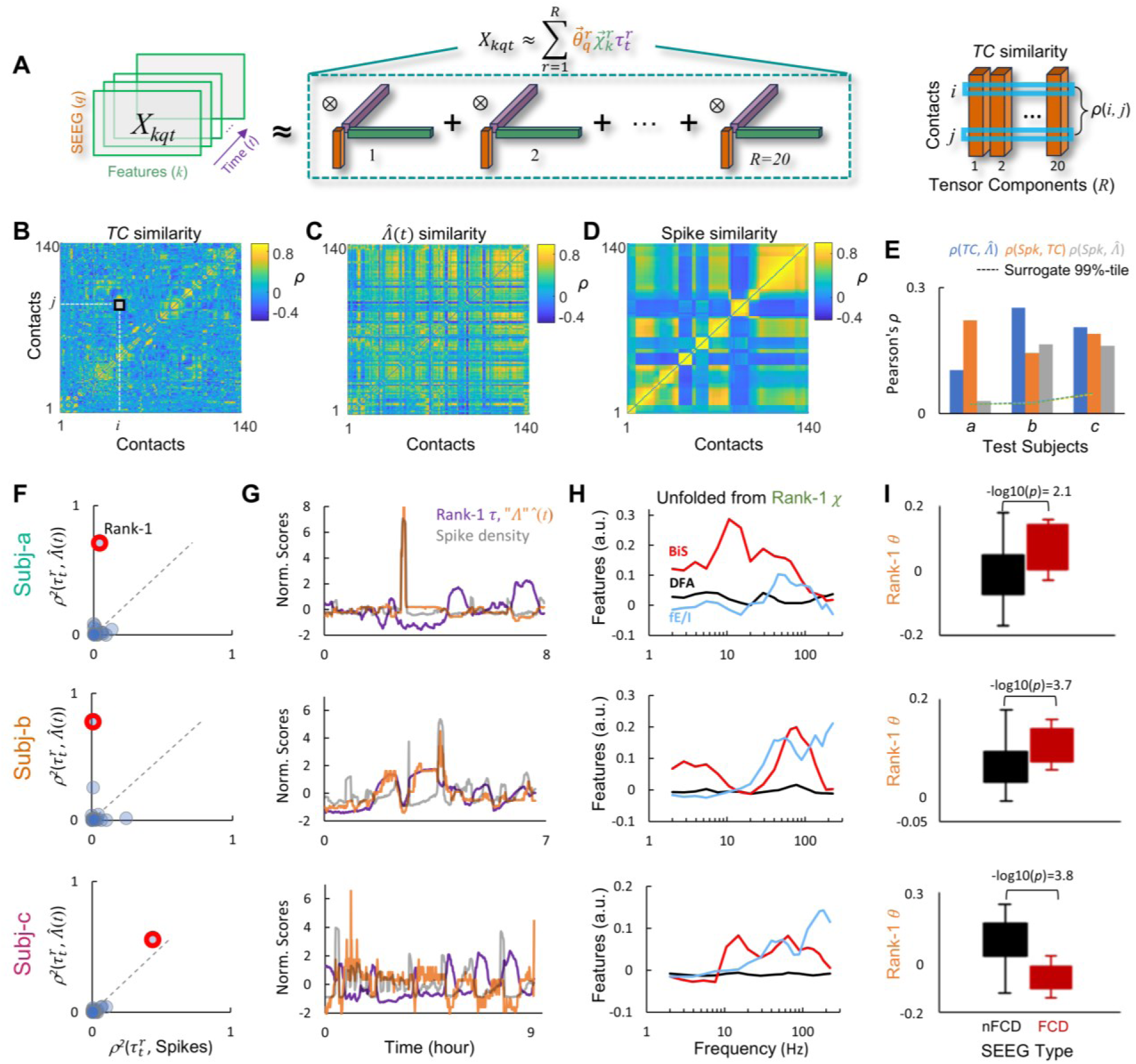
Tensor component (TC) analysis identifies components that correlate with EpiNet-saliency and spikes while revealing individual-specific details. (A) Left: tensor decomposition of one subject’s raw sleep features *X_kqt_*, where, for example in subject *b*, *q* = 140 contacts, each with *k* = 260 raw features, observed over *t* = 400 temporal windows. ⊗: outer product. Right: inter-contact similarity is the correlation between their TCs, with subject *b*’s TC similarity as an example shown in (B). (B–D) An example of one subject’s inter-contact similarity by (B) TC, (C) EpiNet-saliency 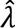(*t*), and (D) spike density. (E) Cross correlations between the three contact similarity matrices for the three test subjects. Dashed: confidence intervals of 1,000 surrogates. (F–I) TC analysis for the three test subjects. (F) Effect size of Spearman’s correlation (*ρ^2^*) between time-loading 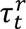 (purple vectors in A) and the global EpiNet-saliency (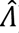) as a function of correlations between the 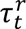 and spikes. Markers: 20 TCs. (G) Time series rank-1 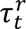 highlighted in (F), global EpiNet-saliency, and spikes. Normalization: (*x* −*mean*(*x*))/*std*(*x*). (H) Visualization of raw feature of the rank-1 feature-vector *χ* (green vector in A). BiS: bistability index; DFA: critical exponent; fE/I: functional E/I index. (I) Differences (unpaired t-test) between contacts inside focal cortical dysplasia (FCD) and non-FCD in the rank-1 contact-vector *θ* (purple vector in A).

where each term 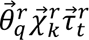 in the sum corresponds to a *r*-th component’s rank-1 approximation of the tensor (we set *R* = 20), with the contact vector 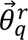 capturing a spatial pattern across *q* contacts showing *k* = 260 raw features captured by the feature vector 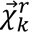, and the temporal profile vector 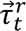 capturing a specific temporal pattern across all time points *t*. Summing over *R* such rank-1 components obtains a rank-*R* approximation of the raw sleep feature tensor *X*_*kqt*_ for one test subject.

## Results

The study was conducted in three steps. We first decomposed 260 raw features from the training dataset into a 10-dimensional eigenfeature space relevant to pathology identified by clinicians. Within this latent space, we trained unsupervised classifiers to construct a parsimonious model for assessing EpiNet-saliency (Methods, Model Design). We next tested the model on three independent subjects (who were not used in model training), using eigenfeatures extracted from 10-minute sliding windows spanning 7–9 hours of sleep-SEEG recordings. Finally, to ensure rigor, we validated the model’s predictions using within-subject tensor component analysis, which jointly captured spatiotemporal feature patterns and assessed their correlation with both EpiNet-saliency and spike density dynamics.

### Raw feature overlap between SZ and nSZ limits supervised classification performance

The EpiNet is hypothesized to exhibit aberrant criticality and strong phase synchrony. For criticality features in narrow-band oscillations, we used DFA to assess long-range temporal correlations (LRTCs), BiS to evaluate bistability, and fE/I to estimate the excitation/inhibition. Synchrony features were derived from narrow-band phase-locking value. Large *first-order* feature estimates characterize densely connected hubs, while large *second-order* feature estimates indicate densely interconnected neighborhoods (Methods). When pooling all 7,183 contacts from the training set, we observed modest to strong correlations among synchrony features (mean *r*^2^ = 0.211) and among criticality features (mean *r*^2^ = 0.114), but weak correlations between criticality and synchrony (mean *r*^2^ = 0.004), indicating little shared information. The *r*^2^ among synchrony features was greater than that among criticality features (unpaired t-test, -log10(*p*) > 12; Supplementary Fig. 2), suggesting that synchrony features are more redundant than criticality features.

During interictal periods, SEEG contacts from clinically identified seizure zone (SZ) typically exhibit pronounced bistable oscillations (Fig. 2D–E), along with larger LRTCs (Fig. 2F) and stronger inhibition (Fig. 2G). Compared to non-SZ contacts (*n* = 5,005), SZ contacts (*n* = 2,178) show a trend toward increased bistability and inhibition in the 15–225 Hz range, slightly elevated LRTCs in the 2–5 Hz and 30–225 Hz bands, and stronger connectivity with a similar spectral profile in the 2–300 Hz range across all first- and second-order synchrony features (Fig. 2I, Supplementary Fig. 3).

To replicate our previous finding on a smaller cohort using one supervised classifer^18^, we trained an additional classifier to localize the SZ. Both classifiers achieved similar accuracy in identifying the SZ (Fig. 2J), with a mean area under the receiver operating characteristic curve (AUC) of 0.74±0.14 (range 0.41–0.98). This variability in classification accuracy suggests substantial overlap between SZ and nSZ contacts. It may reflect the presence of nSZ contacts with SZ-like characteristics or result from inter-subject variability in the differences between SZ and nSZ.

### Differentiate the seizure zone using eigenfeatures

To assess whether epileptogenic features are captured by a manageable number of latent variables for model training (Fig. 3A), we projected raw interictal-SEEG features into the eigenfeature space (Eq. 1–2) to obtain coefficients (*Φ*). We assessed the cohort-level difference between SZ and nSZ contacts in each coefficients using Cohen’s *d*. Coefficients were ranked accordingly (Fig. 3B), with the rank-1 coefficient showing the largest *d* and thus the strongest separation between SZ and nSZ. This cohort analysis was validated by within-subject analysis (gray line, Fig. 3B). In the cohort-pooled samples, over 50 coefficients differentiated SZ from nSZ contacts compared to contact label–shuffled surrogates (*p* < 0.001), with the eigenfeatures of the top three components capturing the most relevant epileptogenic features. The rank-1 criticality eigenfeature closely matches the SZ feature patterns reported previously^18^, showing strong β–γ band bistability and inhibition (top, Fig. 3C), consistent with high-seizure-risk regimes in generative models^15,18^. In contrast, the rank-1 synchrony eigenfeatures are similar across frequencies (bottom, Fig. 3C). Notably, the eigenfeatures of rank-2 and 3 coefficients reflect increased γ and β band excitation, respectively (Supplementary Fig. 1B), suggesting a shift toward a more excitation-dominant regime—one that immediately precedes a transition to seizure-like supercritical hypersynchrony in models^18^.

The joint distribution of the rank-1 and 2 coefficients forms a funnel-like surface, with most contacts concentrated near its base (cool-hot color scale as the z-axis in Fig. 3D). Across subjects (arrows in Fig. 3D), the mean positions of SZ and nSZ contacts were separated, with SZ-vs-nSZ differences quantified by arrow length. Since the coefficients were derived from only 10-minute of interictal SEEG, it remains unclear whether this variability reflects a specific phase within a longer seizure cycle at the time of recording or the mixing of nEZ between the EpiNet and non-EpiNet. Nonetheless, the observed smooth manifold suggests that it is feasible to use the coefficients to train a model to capture time-varying separability between the hypothetical EpiNet and the non-EpiNet.

### Identify the EpiNet using ten eigenfeature coefficients

For simplicity, we used the ten most significant coefficients (Fig. 3B) to classify the EpiNet. Two classifiers were trained in parallel, and their agreement defined the *consensus EpiNet membership* ( *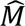* ∈ {0, 1}, Methods), with 1 indicating highest likelihood. Visualizing the average *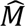* in the joint distribution of rank-1 and rank-2 coefficients revealed a funnel-like surface (with the cool–hot color scale in Fig. 3E representing the z-axis), in which the EpiNet contacts (*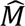* → 1) and the rest are well separated. Notably, the within-subject mean of SZ contacts (arrowheads in Fig. 3D) predominantly fall within the EpiNet region (hot-color in Fig. 3E), whereas the mean of nSZ contacts (arrow-tails in Fig. 3D) are mostly located in the non-EpiNet area (cool-color in Fig. 3E).

We next investigated whether differences in *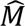* estimates between SZ and nSZ account for variability in supervised SZ-classification accuracy reported previously^18,26^ (Fig. 2I) The within-subject difference in mean *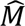* between SZ and nSZ correlates with the mean AUC of the two supervised classifiers for the SZ, with an effect size (*r*^2^) of 0.5 (Fig. 3F). This suggests that when EpiNet nSZ and non-EpiNet nSZ regions are mixed (*i.e.*, EpiNet is more diffused into nSZ areas) or when the entire network is in a low seizure-risk phase (*i.e.*, a 10-minute window is insufficient to capture EpiNet features), the difference between SZ and nSZ in *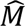* diminishes, leading to reduced supervised classification accuracy.

### Construct a Saliency-Model in a 2-dimensional latent space

To determine whether EpiNet-membership estimates reflect epileptogenicity, we assessed correlation between *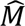* estimates with spike density—a clinical biomarker for seizure risks (Fig. 3G). We found a moderate but significant correlation between *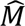* and spike density (Spearman’s *r* = 0.31, -log10(*p*) > 9.15, in total *n* = 371 non-zero pixels from the joint distributions in Fig. 3E&G), suggesting that the EpiNet membership indeed reflects seizure risk.

The EpiNet-saliency model was constructed by fitting the joint distribution of *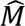* as a smooth surface in the (*Φ_1_–Φ_2_*) coordinates representing saliency assessment 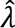 ∈ {0, 1}, where 1 denotes highest saliency. A parsimonious EpiNet-saliency model can be implemented as a lookup table, using the rank-1 and 2 coefficients from new data as inputs to obtain a saliency assessment, thereby reducing feature space to 0.77% of its original size and eliminating the need for re-training classifiers for new data.

### Test the EpiNet-saliency model in an independent cohort

To demonstrate the potential clinical value, we tested the model on three novel patients diagnosed with type-2 focal cortical dysplasia (FCD). FCD was selected because it serves as a prototypical clinical model of sleep-related epilepsy with a well-characterized histopathological substrate^42^. All three patients achieved seizure freedom (Engel class 1A) at a minimum of one-year follow-up, confirming accurate EZ localization in presurgical workup and following resection. From subject *a* to *c*, clinical seizure risk increased, as reflected by a rising number of observed seizures during pre-surgical monitoring and a greater number of contacts located in dysplastic cortex, confirmed by post-surgical histopathological analysis (Fig. 4B). Raw features were extracted for 10-minute sliding window with one-minute step size from 7–9 hours of sleep recorded over one night, enabling us to examine ultradian fluctuations in EpiNet-saliency as in previous studies^27–29^.

We projected the time-resolved sleep-SEEG features into the eigenfeature space (Fig. 1E) to obtain the rank-1 and rank-2 coefficients (*Φ_1_* and *Φ_2_*). As a sanity check before model testing, we visualized the probability distributions of raw features and coefficients for both the training data and test data. The sleep-SEEG distributions closely resembled those of interictal-SEEG and showed no prominent outliers (Fig. 4C), indicating strong similarity between datasets.

Slow fluctuations in the time-resolved *Φ_1_* and *Φ_2_* were observed in all three subjects (an example is shown in Fig. 4D). In the *Φ_1_*–*Φ_2_* state space, the temporal trajectories of FCD and non-FCD contacts are separated (Fig. 4E). This separation becomes more apparent when time-averaging the *Φ_1_* and *Φ_2_* estimates for each contact (Fig. 4F). This distinction between epileptogenic FCD and non-FCD contacts is consistent with the separation observed between SZ and nEZ contacts in the training set (Fig. 3D), both supporting the proposed state space (Fig. 1C). Notably, abrupt changes in both coefficients coincided with sudden increases in time-resolved spike density (Fig. 4G), suggesting that these coefficients likely capture slow fluctuations in epileptogenicity.

### Dynamical relationship between spike density and EpiNet-saliency

To investigate the correlation between EpiNet-saliency estimates and spike occurrences over time, we computed the mean spike density across contacts and applied the Hilbert transform to extract phase time series of the global spike density (middle, Fig. 4G). In all test subjects, FCD contacts showed larger saliency assessments across all spike phases (top row, Fig. 4H), with effect sizes (bottom row, Fig. 4H), suggesting a trend of positive correlation with clinically identified seizure risk (Fig. 4B).

To better assess this trend, we grouped all samples into four 90° spike phase bins, revealing an unexpected dichotomy in spike-related EpiNet-saliency modulation. As global spike density increased (0–90°), saliency in FCD contacts decreased in subjects *a* and *b* but increased in subject *c* who had the highest saliency (top row, Fig. 4I). When spike density decreased (90– 180°), saliency in FCD contacts continued to decline in subject *a* and *b* but kept rising in *c*.

Global EpiNet-saliency, defined as the average across all contacts, varied over time (Fig. 4J, top), corroborates with fluctuations in classification accuracy based on contact-wise saliency (Fig. 4J, bottom). Peak AUC of the classification reached 0.63, 0.85, and 0.94 in three test subjects. Both global and local EpiNet-saliency were consistent with clinical assessments of seizure risk, revealing a clear gradient of increasing risk from subject *a* to *c*. These results validate the model and offer new support for the low-dimensional epileptogenicity hypothesis at both local and global levels.

### Tensor component analysis validates EpiNet-saliency Assessments

While the model provides efficient but coarse-grained assessments of clinically relevant EpiNet-saliency, it remains unclear whether it overlooks subject-specific details that may be clinically important—or whether the time-varying saliency estimates truly reflect subject level dynamics. To address this, we factorized each test subject’s sleep feature *X*_*kqt*_, a three-way tensor, into 20 tensor components (TC), where each TC represents a latent feature as the outer product of three vectors (Fig. 5A)—the contact vector 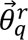 quantifies how strongly each contact expresses a raw feature pattern characterized by the feature vector 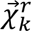 representing a raw feature pattern, which evolves over time according to the time-course vector 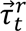.

We first assessed the spatial similarity between the TCs, EpiNet, and spike density. All-to-all correlations were computed between contacts using their TC weights, the time course of EpiNet-saliency, and spike density. In a TC similarity matrix, a strong Pearson correlation between two contacts (*i*, *j*) across all contact vectors indicates that they share similar raw feature patterns and temporal dynamics across all components (Fig. 5A). Compared to saliency- and spike-derived similarity, the TC matrix reveals more detailed information in contact interactions (Fig. 5B–D). These matrices were moderately correlated across subjects (Fig. 5E), suggesting that they capture shared aspects of the underlying pathological dynamics.

Next, we examined whether the TC time courses correlated with global spike density (*e.g.*, blue trace in Fig. 4G, middle panel) and global EpiNet-saliency (Fig. 4J, top panel). In subject *a* and *b*, only one out of the 20 TCs showed correlation between its time-course and EpiNet-saliency, but not with spike density (Fig. 5F–G). In subject *c*, one TC correlated with both saliency and spike density. These findings independently validated model’s EpiNet saliency assessment.

We then analyzed the generalizability and specificity of the raw feature patterns in the tensor component most strongly associated with EpiNet-saliency (*i.e.*, the rank-1 TC highlighted in Fig. 5F). The corresponding feature vector showed moderate inter-subject similarity, with Pearson’s correlations of *r*(*a*, *b*) = 0.23, *r*(*a*, *c*) = 0.21, and *r*(*b*, *c*) = 0.77. Notably, subject *b* and *c* are characterized by elevated bistability in 30–200 Hz and 10–200 Hz, consistent with previous findings^15,18^, whereas *a*, who had the fewest seizures during presurgical monitoring, showed elevated bistability centered at 10 Hz but not in γ band (Fig. 5H, blue line). All three test subjects exhibited increased excitation, most prominently above 30 Hz, which contrasts with increased inhibition in the same band in the training subjects (Fig. 3C, blue line).

Lastly, we compared FCD and non-FCD contacts based on the contact vector 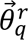 of the rank-1 EpiNet-relevant TC, where higher values indicate stronger expression of the component (Fig. 5H). In subject *a* and *b*, the FCD contacts showed higher values than non-FCD contacts (Fig. 5I) with large effect sizes (Cohen’s *d* = −0.99 and −1.1, corresponding to 2.7 and 3.6 standard deviations (SD) above the mean of 10^4^ surrogates, respectively). In subject *c*, however, the non-FCD contacts showed greater values (Cohen’s *d* = 1.3, 3.8 SD above the surrogate mean).

In summary, the TC analysis uncovered subject-specific details not captured by the EpiNet-saliency model and suggests that fluctuations in EpiNet-saliency are sparsely encoded within individual tensor components.

## Discussion

We aimed to address a long-standing challenge in understanding the systems-level mechanisms of epilepsy^3,43,44^. We asked whether the EpiNet can be characterized by invariant landmarks in a low-dimensional latent space derived from hundreds of neuronal features of interictal SEEG data. This hypothesis is inspired by bifurcation theory, which describes local seizure dynamics using only four canonical bifurcations^45,46^. We combined dimensionality reduction and interpretable classifiers to train a model to assess EpiNet-saliency in a two-dimensional eigenfeature space and cross-domain tested the model. The model not only identified clinically relevant EpiNet-saliency but also revealed unexpected, subject-specific relationships between the saliency and interictal spikes. Independent tensor component analysis further validated the model, reinforcing its rigor.

EpiNet-saliency assessments were positively correlated with spikes in both training and test subjects, and therefore likely reflects seizure risk or proximity to seizure onset, as proposed in previous studies^39,45^. However, since only interictal data were available for training in this study, the exact locations of seizure bifurcation—onset, offset, and refractory state—were not mapped within the eigenfeature space. Therefore, it is important to emphasize that EpiNet-saliency in the present work should be interpreted as reflecting a degree of epileptogenic proneness, rather than as a measure of distance to bifurcation or a predictor of seizure imminence.

### Novelty: a framework for unified biomarkers

While mechanistic understanding of the EZ has advanced significantly over the last seven decades^43^, presurgical EZ-localization remains a complex process relying on multidisciplinary expertise and extensive clinical infrastructures^3,9,47^. The sensitivity and specificity of any single biomarker is nonetheless limited by various conceptual and practical factors^8,9,39,44,48–50^, and no existing framework is capable of comprehensively utilizing all relevant biomarkers.

To fill this gap, we recently proposed shifting the focus from a discrete EZ to a broader epileptogenic network (EpiNet), combining multiple biomarkers with machine-learning to achieve more comprehensive localization^18,26^. The current study aimed to resolve the high-dimensionality challenge inherent in our earlier approach and provided initial proof of concept for a scalable framework to decipher systems-level complexity of the EpiNet, bridging novel biomarkers and data science approaches with clinical needs.

### EpiNet characterized in a latent space

The model takes two eigenfeatures as input, reducing dimensionality to 0.77% of the original features—without compromising individual sensitivity due to band-wise feature averaging in previous studies^15,18^. This not only resolves the challenge in model training but simplifies interpretation. The rank-1 eigenfeature unfolds as elevated inhibition, bistability, and LRTCs in 30–225 Hz oscillations, indicative of aberrant criticality with a catastrophic shift—a pattern in line with findings from both theoretical and empirical studies^15,18,22^. The EpiNet-saliency in new data can be assessed by projecting the raw features into the eigenfeature space and retrieving the saliency score from the trained model, eliminating the need for retraining. This approach significantly reduces computational overhead, supports analysis of large clinical datasets, and may potentially be adapted for online seizure forecasting.

### EpiNet characterized on a minute-level timescale

Seizure prediction is most meaningful on a minute-level timescale, as it not only allows sufficient time for intervention^51,52^, improves computational efficiency^26^, but also grounded in physiological foundations. Bifurcation models^39,45,46,53^ suggest a slow *permittivity* variable that controls an ensemble’s operating distance to seizure bifurcation. Numerous mechanisms—such as synaptic, metabolic, electrophysiological, and anatomical/functional changes—may synergize and contribute to temporal fluctuations in the permittivity. This suggests that the seizure dynamics should not be attributed to a few isolated mechanisms, but rather treated as a latent variable^45^, as considered here.

In the sleep-SEEG, global EpiNet-saliency fluctuated on a multi-minute scale and correlated with spike density and rank-1 tensor component. At the local level, SZ and nSZ exhibited time-varying contrast in EpiNet-saliency, reflecting ultradian evolution in seizure risks^27–29^. Similar fluctuations are likely present in the interictal-SEEG subjects. As a result, the SZ-vs-nSZ contrast during 10-minute interictal-SEEG could have varied substantially across subjects. This variability would lead to poor classification accuracy, particularly for subjects recorded during a low seizure risk phase when the contrast is weak. This individual variability, however, is desired in model training as it would generalize to the gradient in new data.

### Large individual variability in EpiNet features

The finding that many non-SZ contacts resemble SZ features suggests that even physiologically normal regions can operate in a pathological regime as previously suggeseted^18^. Clinically, this reinforces the notion that some regions may facilitate seizure without being able to independently generate seizures. Such regions may be suboptimal targets for resection but suitable candidates for neuromodulation^9,52,54^.

All three test subjects showed considerable fluctuations in EpiNet-saliency during sleep, while their bistability profiles—derived from the rank-1 EpiNet-relevant tensor component—differed significantly, despite sharing the same FCD histopathology. One patient exhibited broad α–γ band bistability and experienced the most frequent seizures, while another showed only α-band bistability and had only one observed seizure.

Collectively, these findings suggest that although functional biomarkers offer greater granularity than structural pathology alone^3,9^, the timing of SEEG recording is crucial. If data are only sampled during a low-risk phase, the EpiNet extent may be underestimated, and its feature pattern may also generalize poorly. This dynamic property of EpiNet-saliency is both a strength and a limitation—the temporal sampling bias warrants future investigation.

### Inhibition and excitation coexist in the latent space

Recent computational modelling has shown that in a critical-like regime with high bistability, increasing neuronal coupling is associated with local dynamics to first exhibit inhibition, then excitation, and eventually supercritical, seizure-like hypersynchrony^18^. The SZ in both interictal- and sleep-SEEG showed elevated bistability, implying a regime shift toward supercritical regime^15,18,21^.

In the rank-1 eigenfeature of interictal-SEEG, increased 30–225 Hz inhibition was observed, whereas increased 30–225 Hz excitation characterized both the sleep-SEEG and lower-rank eigenfeatures of the interictal-SEEG. This suggests that some interictal-SEEG subjects could have been closer to seizure onset than others. Given that FCD is frequently associated with sleep-related epilepsy^42^, it is possible that the test subjects were indeed closer to seizure onset during sleep than most of the interictal-SEEG subjects—likely operating on the overexcited, supercritical side of the critical regime as our modelling study has suggested^14^. Despite this potential between-cohort difference, the model cross-domain identifies the FCD contacts, suggesting that variation in a specific raw feature set had little impact on overall prediction.

### A potential dichotomy of excitatory and inhibitory spikes

Interictal spikes, albeit an established clinical biomarker, often appear across widespread brain regions outside seizure foci, diminishing their specificity^55,56^. To address this variability, our time-resolved sleep-SEEG analysis revealed two distinct correlation patterns between spikes and EpiNet-saliency. In subject *a*, saliency increased following a rise in global spike and peaked at the trough of spike activity. In contrast, the other two subjects exhibited the opposite pattern. Notably, subject *a*, who experienced most frequent seizures during presurgical evaluation, was the only case whose pathological tensor component positively correlated with both EpiNet-saliency and spikes. This individual variability may explain the modest correlation observed in the training data (*r* = 0.31), as mixing patients with opposing patterns may obscure trends. These finding support the hypothesis of a functional dichotomy of interictal spikes: some may be excitatory and pro-epileptogenic while others are inhibitory and protective^8,57–62^. Therefore, spikes should not be used indiscriminately as a biomarker.

### Limitations and future efforts

This innovative approach facilitates the inclusion of additional biomarkers and, importantly, may provide an answer to why established clinical biomarkers often lose specificity^25^. However, it remains unclear whether histopathological substrates (*e.g.*, hippocampal sclerosis vs FCD), comorbidities, or brain states are encoded in separate latent variables or interact with those relevant to the EpiNet. Such interactions may exist in the sleep-SEEG, as ultradian fluctuations in seizure risk is likely driven by the NREM cycles, where epileptiform activity is typically suppressed during REM sleep and promoted during stage-2 NREM sleep^27,63^, likely over large-scale connectivity^22^. Comorbidities are prevalent among epilepsy patients^64^, but due to limited clinical information we were unable to investigate them here.

Our model was trained on interictal SEEG data. As such, EpiNet-saliency should be interpreted as the likelihood that a region is part of the EpiNet based what the model learned from systems-level features—rather than as a direct measure of proximity to seizure bifurcation, as in models^45^ and SEEG studies^39^ based on local dynamics. Seizure onset can be classified using electrophysiological signatures generated by four known bifurcations^39,45,53^. Therefore, it is feasible to empirically estimate the distance to bifurcation with the biomarkers employed here and in recent studies^22,24,65^—but at a second-level resolution, rather than minute-level resolution considered here. Despite its gold-standard signal quality^47^, SEEG is costly, invasive, limited in accessibility, making it unsuitable for repeated measures or large-scale screening. SEEG implants are guided by clinical hypotheses, which may lead to under-sampling in complex cases^66^. As a next step, we will test this approach for whole-brain, non-invasive mapping with source-reconstructed MEG or scalp EEG^67,68^.

## Conclusion

This work advances a methodological breakthrough for hypothesis-free characterizing the EpiNet. It sets a clear line of sight from this innovative approach through a powerful framework that synergizes information across diverse biomarkers—aimed at advancing collaborative standardization in presurgical EZ mapping, mechanistic studies, and beyond^69^.

## Data availability

Compliance with Italian governing laws and Ethical Committee regulations prohibits the sharing of raw SEEG data and patient information. However, interim data and final results that support the findings of this study can be obtained from the corresponding authors upon reasonable request. The python toolbox for feature extraction can be obtained from the corresponding authors upon reasonable request.

## Acknowledgement

This study has been supported by a Sigrid Jusélius Foundation Fellowship awarded to SHW (210527). LN and GA were supported by NEXTGENERATIONEU (NGEU) and by the Ministry of University and Research (MUR), National Recovery and Resilience Plan (NRRP), project MNESYS (PE0000006)—A Multiscale integrated approach to the study of the nervous system in health and disease (DN. 1553 11.10.2022). This project was also funded by: Academy of Finland grant (SA 253130 and 296304) awarded to JMP and Sigrid Jusélius Foundation grant awarded to SP and JMP. MM was supported by the DARLING Project (Projet-ANR-19-CE48-0002) obtained by PC from Agence Nationale de la Recherche, France. SHW was partially supported by the DARLING Project.

## Author contributions

Funding acquisition: SHW, JMP, SP, PC

Supervision: JMP, PC Conceptualization: SHW

Methodology: SHW, MM, GA, JMP

Resource: LN, GA

Software: SHW, GA, VM

Formal analysis: SHW, MM, VM

Visualization: SHW, PF, JMP

Writing - Original Draft: SHW, PF, LN, JMP

All authors edited and approved the final draft.

## Abbreviations

AUC: area under the receiver-operating characteristic curve
BiS: bistability index
DFA: the exponent obtained using detrended fluctuation analysis
DRE: drug-resistant epilepsy
EpiNet: epileptogenic network
EZ: epileptogenic zone
FCD: focal cortical dysplasia
fE/I: functional E/I balance index
LRTCs: long-range temporal correlations
PLV: phase-locking value
SEEG: stereo-electroencephalography
TC: tensor component
SVD: singular value decomposition
SZ: seizure zone

## References

1. Devinsky O, Vezzani A, O’Brien TJ, et al. Epilepsy. Nat Rev Dis Prim. 2018;4(May). doi:10.1038/nrdp.2018.24

2. Luders H, Engel J, Munari C. Surgical Treatment of the Epilepsies. Raven Press; 1993.

3. Zijlmans M, Zweiphenning W, van Klink N. Changing concepts in presurgical assessment for epilepsy surgery. Nat Rev Neurol. 2019;15(10):594–606. doi:10.1038/s41582-019-0224-y

4. Stacey W, Kramer M, Gunnarsdottir K, et al. Emerging roles of network analysis for epilepsy. Epilepsy Res. 2020;159(December 2019):106255. doi:10.1016/j.eplepsyres.2019.106255

5. Khambhati AN, Davis KA, Oommen BS, et al. Dynamic Network Drivers of Seizure Generation, Propagation and Termination in Human Neocortical Epilepsy. PLoS Comput Biol. 2015;11(12):1–19. doi:10.1371/journal.pcbi.1004608

6. van Lanen RHGJ, Colon AJ, Wiggins CJ, et al. Ultra-high field magnetic resonance imaging in human epilepsy: A systematic review. NeuroImage Clin. 2021;30(February):102602. doi:10.1016/j.nicl.2021.102602

7. Jacobs J, Zijlmans M, Zelmann R, et al. High-frequency electroencephalographic oscillations correlate with outcome of epilepsy surgery. Ann Neurol. 2010;67(2):209–220. doi:10.1002/ana.21847

8. Guth TA, Kunz L, Brandt A, et al. Interictal spikes with and without high-frequency oscillation have different single-neuron correlates. Brain. 2021;144(10):3078–3088. doi:10.1093/brain/awab288

9. Miller KJ, Fine AL. Decision-making in stereotactic epilepsy surgery. Epilepsia. Published online November 1, 2022. doi:10.1111/epi.17381

10. Astner-Rohracher A, Zimmermann G, Avigdor T, et al. Development and Validation of the 5-SENSE Score to Predict Focality of the Seizure-Onset Zone as Assessed by Stereoelectroencephalography. JAMA Neurol. 2022;79(1):70–79. doi:10.1001/jamaneurol.2021.4405

11. Cardinale F, Rizzi M, Vignati E, et al. Stereoelectroencephalography: Retrospective analysis of 742 procedures in a single centre. Brain. 2019;142(9):2688–2704. doi:10.1093/brain/awz196

12. Bulacio JC, Bena J, Suwanpakdee P, et al. Determinants of seizure outcome after resective surgery following stereoelectroencephalography. J Neurosurg. 2022;136(6):1638–1646. doi:10.3171/2021.6.JNS204413

13. Auno S, Lauronen L, Wilenius J, Peltola M, Vanhatalo S, Palva JM. Detrended fluctuation analysis in the presurgical evaluation of parietal lobe epilepsy patients. Clin Neurophysiol. 2021;132(7):1515–1525. doi:10.1016/j.clinph.2021.03.041

14. Fuscà M, Siebenhühner F, Wang SH, et al. Brain criticality predicts individual synchronization levels in humans. Nat Commun. Published online 2023:2022.11.24.517800. 10.1038/s41467-023-40056-9

15. Wang SH, Siebenhühner F, Arnulfo G, et al. Critical-like brain dynamics in a continuum from second-to first-order phase transition. J Neurosci. 2023;43(45):7642–7656. doi:10.1523/JNEUROSCI.1889-22.2023

16. Beggs JM. The criticality hypothesis: How local cortical networks might optimize information processing. Philos Trans R Soc A Math Phys Eng Sci. 2008;366(1864):329-343. doi:10.1098/rsta.2007.2092

17. Monto S, Vanhatalo S, Holmes MD, Palva JM. Epileptogenic neocortical networks are revealed by abnormal temporal dynamics in seizure-free subdural EEG. Cereb Cortex. 2007;17(6):1386–1393. doi:10.1093/cercor/bhl049

18. Wang SH, Arnulfo G, Nobili L, et al. Neuronal Synchrony and Critical Bistability: Mechanistic Biomarkers for Localizing the Epileptogenic Network. Epilepsia. Published online 2024. 10.1101/2023.05.21.541570

19. van van Hugte EJH, Schubert D, Nadif Kasri N. Excitatory/inhibitory balance in epilepsies and neurodevelopmental disorders: Depolarizing γ-aminobutyric acid as a common mechanism. Epilepsia. 2023;64(8):1975–1990. doi:10.1111/epi.17651

20. Lepeu G, van Maren E, Slabeva K, et al. The critical dynamics of hippocampal seizures. Nat Commun. 2024;accepted(July 2023). doi:10.1038/s41467-024-50504-9

21. Thom R. Structural Stability and Morphogenesis. 1st Ed. CRC Press; 1972. https://www.taylorfrancis.com/books/9780429493027

22. Burlando G, Belforte C, Siebenhühner F, et al. Sleep-Modulated Cross-Frequency Coupling Between δ Phase and β-γ Bistability: A System-Level Modulation of Epileptic Activity. bioRxiv. 2025;(125):1–30. https://www.biorxiv.org/content/10.1101/2025.03.25.642592v1

23. Siebenhühner F, Wang SH, Arnulfo G, et al. Genuine cross-frequency coupling networks in human resting-state electrophysiological recordings. Poeppel D, ed. PLOS Biol. 2020;18(5):e3000685. doi:10.1371/journal.pbio.3000685

24. Myrov V, Siebenhühner F, Juvonen JJ, Arnulfo G, Palva S, Palva JM. Rhythmicity of neuronal oscillations delineates their cortical and spectral architecture. Commun Biol. 2024;7(1):1–18. doi:10.1038/s42003-024-06083-y

25. Zweiphenning W, van MA, C Klink NE, et al. Intraoperative Electrocorticography Using High-Frequency Oscillations or Spikes to Tailor Epilepsy Surgery in the Netherlands (the HFO Trial): A Randomised, Single-Blind, Adaptive Non-Inferiority Trial. Vol 21.; 2022. www.thelancet.com/neurology

26. Wang S, Marzulli M, Arnulfo G, et al. Machine learning models trained in a low-dimensional latent space for epileptogenic zone (EZ) localization. In: EUSIPCO.; 2024.

27. Karoly PJ, Rao VR, Gregg NM, et al. Cycles in epilepsy. Nat Rev Neurol. 2021;17(5):267-284. doi:10.1038/s41582-021-00464-1

28. Baud MO, Kleen JK, Mirro EA, et al. Multi-day rhythms modulate seizure risk in epilepsy. Nat Commun. 2018;9(1):1–10. doi:10.1038/s41467-017-02577-y

29. Leguia MG, Andrzejak RG, Rummel C, et al. Seizure Cycles in Focal Epilepsy. JAMA Neurol. 2021;78(4):454–463. doi:10.1001/jamaneurol.2020.5370

30. Arnulfo G, Wang SH, Myrov V, et al. Long-range phase synchronization of high-frequency oscillations in human cortex. Nat Commun. 2020;11(1). doi:10.1038/s41467-020-18975-8

31. Arnulfo G, Hirvonen J, Nobili L, Palva S, Palva JM. Phase and amplitude correlations in resting-state activity in human stereotactical EEG recordings. Neuroimage. 2015;(112):114–127.

32. Linkenkaer-Hansen K, Nikouline V V., Palva JM, Ilmoniemi RJ. Long-Range Temporal Correlations and Scaling Behavior in Human Brain Oscillations. J Neurosci. 2001;21(4):1370–1377. doi:10.1523/jneurosci.21-04-01370.2001

33. Bruining H, Hardstone R, Juarez-Martinez EL, et al. Measurement of excitation-inhibition ratio in autism spectrum disorder using critical brain dynamics. Sci Rep. 2020;10(1). doi:10.1038/s41598-020-65500-4

34. Diachenkoa M, Sharma A, Smit DJA, et al. Functional excitation-inhibition ratio indicates near-critical oscillations across frequencies. Imaging Neurosci. Published online 2024.

35. Freyer F, Roberts JA, Ritter P, Breakspear M. A Canonical Model of Multistability and Scale-Invariance in Biological Systems. PLoS Comput Biol. Published online 2012. doi:10.1371/journal.pcbi.1002634

36. Breiman L. Random Forests. In: Machine Learning. Vol 45. Kluwer Academic Publishers; 2001:5-32.

37. Chen T, Guestrin C. XGBoost: A scalable tree boosting system. Proc ACM SIGKDD Int Conf Knowl Discov Data Min. 2016;13–17-Augu:785-794. doi:10.1145/2939672.2939785

38. Pedregosa F, Varoquaux G, Gramfort A, Michel V, Thirion B. Scikit-learn: Machine Learning in Python. J Mach Learn Res. 2011;12:2825–2830. doi:10.1289/EHP4713

39. Saggio ML, Crisp D, Scott JM, et al. A taxonomy of seizure dynamotypes. Elife. 2020;9:1–56. doi:10.7554/eLife.55632

40. Warner J, The scikit-fuzzy development team. scikit-fuzzy: Fuzzy Logic Toolbox for Python. Published 2018. github.com/scikit-fuzzy/scikit-fuzzy

41. Kolda TG, Bader BW. Tensor decompositions and applications. SIAM Rev. 2009;51(3):455-500. doi:10.1137/07070111X

42. Gibbs SA, Proserpio P, Francione S, et al. Clinical features of sleep—related hypermotor epilepsy in relation to the seizure—onset zone: A review of 135 surgically treated cases. Epilepsia. 2019;(April):707–717. doi:10.1111/epi.14690

43. Lara Jehi. The Epileptogenic Zone: Concept and Definition. Epilepsy Curr. 2018;18(1):12–16.

44. Fisher RS, Scharfman HE, DeCurtis M. How Can We Identify Ictal and Interictal Abnormal Activity? Adv Exp Med Biol. Published online 2014:3–23. doi:10.1007/978-94-017-8914-1_1

45. Jirsa VK, Stacey WC, Quilichini PP, Ivanov AI, Bernard C. On the nature of seizure dynamics. Brain. Published online 2014. doi:10.1093/brain/awu133

46. Saggio ML, Spiegler A, Bernard C, Jirsa VK. Fast–Slow Bursters in the Unfolding of a High Codimension Singularity and the Ultra-slow Transitions of Classes. J Math Neurosci. 2017;7(1):1–47. doi:10.1186/s13408-017-0050-8

47. Frauscher B, Mansilla D, Abdallah C, et al. Learn how to interpret and use intracranial EEG findings. Epileptic Disord. 2024;26(1):1–59. doi:10.1002/epd2.20190

48. Perucca P, Gotman J. Delineating the epileptogenic zone: spikes versus oscillations. Lancet Neurol. 2022;21(11):949–951. doi:10.1016/S1474-4422(22)00396-9

49. Gliske S V., Irwin ZT, Chestek C, et al. Variability in the location of high frequency oscillations during prolonged intracranial EEG recordings. Nat Commun. Published online 2018. doi:10.1038/s41467-018-04549-2

50. Fisher RS, Cross JH, French JA, et al. Operational classification of seizure types by the International League Against Epilepsy: Position Paper of the ILAE Commission for Classification and Terminology. Epilepsia. 2017;58(4):522–530. doi:10.1111/epi.13670

51. Qi Z, Liu H, Jin F, et al. A wearable repetitive transcranial magnetic stimulation device. Nat Commun. 2025;(September 2024):1–14. doi:10.1038/s41467-025-58095-9

52. Nair DR, Laxer KD, Weber PB, Murro AM, Park YD. Nine-year prospective e ffi cacy and safety of brain-responsive neurostimulation for focal epilepsy. Neurology. 2020;0(95(9)). doi:10.1212/WNL.0000000000010154

53. Izhikevich EM. Neural excitability, spiking and bursting. Int J Bifurcat Chaos. 2000;10(6):1171-1266. doi:10.1142/S0218127400000840

54. Rao VR, Rolston JD. Unearthing the mechanisms of responsive neurostimulation for epilepsy. Commun Med. 2023;3(1):1–6. doi:10.1038/s43856-023-00401-x

55. Ung H, Cazares C, Nanivadekar A, et al. Interictal epileptiform activity outside the seizure onset zone impacts cognition. Brain. Published online 2017. doi:10.1093/awx178

56. Smith EH, Merricks EM, Davis T, et al. Human interictal epileptiform discharges are bidirectional traveling waves echoing ictal discharges. Elife. 2022;11(73541):1–20.

57. Shi W, Shaw D, Walsh KG, et al. Spike ripples localize the epileptogenic zone best: an international intracranial study. Brain. 2024;147(7):2496–2506. doi:10.1093/brain/awae037

58. Rasmussen T. Characteristics of a Pure Culture of Frontal Lobe Epilepsy. Epilepsia. 1983;24(4):482–493. doi:10.1111/j.1528-1157.1983.tb04919.x

59. Serafini R. Similarities and differences between the interictal epileptiform discharges of green-spikes and red-spikes zones of human neocortex. Clin Neurophysiol. 2019;130(3):396–405. doi:10.1016/j.clinph.2018.12.011

60. Schaft E V., Sun D, van ‘t Klooster MA, et al. Spatial and temporal properties of intra-operatively recorded spikes and high frequency oscillations in focal cortical dysplasia. Clin Neurophysiol. 2024;162:210–218. doi:10.1016/j.clinph.2024.03.038

61. Karoly PJ, Freestone DR, Boston R, et al. Interictal spikes and epileptic seizures: Their relationship and underlying rhythmicity. Brain. 2016;139(4):1066–1078. doi:10.1093/brain/aww019

62. Curtis M De, Avanzini G. Interictal spikes in focal epileptogenesis. Prog Neurobiol. 2001;63:541–567.

63. Nobili L, Frauscher B, Eriksson S, et al. Sleep and epilepsy: A snapshot of knowledge and future research lines. J Sleep Res. 2022;31(4):1–11. doi:10.1111/jsr.13622

64. Keezer MR, Sisodiya SM, Sander JW. Comorbidities of epilepsy: Current concepts and future perspectives. Lancet Neurol. 2016;15(1):106–115. doi:10.1016/S1474-4422(15)00225-2

65. Dumeur M, Wang SH, Palva JM, Ciuciu P. Multifractality in critical neural field dynamics. Published online 2023:1–6. http://arxiv.org/abs/2312.03219

66. Jaber K, Avigdor T, Mansilla D, et al. A spatial perturbation framework to validate implantation of the epileptogenic zone. Nat Commun. 2024;15(1):1–17. doi:10.1038/s41467-024-49470-z

67. Bagic A, Funke ME, Ebersole J. Magnetoencephalography (MEG)/Magnetic Source Imaging (MSI) in Noninvasive Presurgical Evaluation of Patients With Medically Intractable Localization-related Epilepsy. J Clin Neurophysiol. 2009;26(4):290–293.

68. Maldjian JA, Lee R, Jordan J, et al. ACR White Paper on Magnetoencephalography and Magnetic Source Imaging: A Report from the ACR Commission on Neuroradiology. Am J Neuroradiol. 2022;43(12):E46–E53. doi:10.3174/ajnr.A7714

69. McKee HR, Vidaurre J, Clarke D, et al. It’s About Time! Timing in Epilepsy Evaluation and Treatment. Epilepsy Curr. Published online 2024. doi:10.1177/15357597241238072

