## Supplementary Materials for "Toward Unified Biomarkers for Focal Epilepsy"

By Wang SH et al.

### Supplementary Methods

#### Local criticality assessments

Narrow-band frequency amplitudes were obtained by convolving the broadband data with Morlet wavelets (*m* = 5), spanning 2 to 225 Hz with equal spacing on a *log*_10_ scale. We then assessed long-range temporal correlations (LRTCs), functional excitation/inhibition (E/I), and bistability from the resulting narrow-band signals.

##### Estimate LRTCs

LRTCs is assessed for narrow-band oscillation amplitudes using linear detrended fluctuation analysis (DFA), we refer readers to the paper by Hardstone et al.^1^ for theoretical background and implementation details. Briefly, the DFA exponent quantifies finite-size power-law scaling in amplitude fluctuations of narrow-band signals. It assumes that the gradual evolution of a monofractal time series produces a normal distribution of fluctuations, which can be characterized by second-order statistical moments. In practice, DFA estimates how fast the root mean square (RMS) of local detrended fluctuations, *F*(*L*), grows with increasing window size *L*:

$F(L)=C\cdot L^{\beta}$	(1)

where *C* is a constant, *β* is the DFA exponent. A DFA exponent greater than 0.6 is considered indicative of critical-like dynamics, significantly exceeding the exponent for a random walk (*β*=0.5, see Fig. 2 for fitting examples). To compute DFA for narrow-band amplitude signals, we used 40 temporal windows ranging in size from 20 oscillatory cycles (*e.g*., 2 seconds for a 10 Hz signal) to 2.5 minutes, spaced evenly on a logarithmic scale. For each window size *L_i_*, *F*(*L_i_*) was computed as the mean RMS of the detrended fluctuations, using a 50% overlap between adjacent windows. Windows with more than 10% samples containing interictal epileptiform spikes were excluded from the mean. Linear regression was performed on the log–log plot of *F*(*L*) vs *L* using bi-square fitting, weighted by the square root of the number of observed windows for each window size.

DFA considers oscillation envelopes as proxies for the rate-of-change of neural activity, which is integrated to assess scale-free long-range temporal correlations (LRTCs) in the detrended fluctuations—effectively capturing fluctuations in the “work” of a neuronal oscillator across temporal scales. LRTCs is theoretically and empirically linked to large-scale functional connectivity^2^ and neuronal avalanche dynamics^3,4^. Significant LRCTs (DFA > 0.6) is often used to demarcate the critical regime^5,6^.

##### Estimate functional E/I

We assessed functional excitation/inhibition in narrow-band oscillation amplitudes using the fE/I index^5^:

$fE/I=1-r(w_{amp}(t),w_{nF}(t) )$	(2)

where $w_{amp}(t)$ is the windowed oscillation amplitudes; $w_{nF}(t)$ is the windowed detrend function of the amplitude-normalized narrow-band signal; $r()$ is the Pearson’s correlation between the $w_{amp}(t)$and $w_{nF}(t)$observations. Within the critical regime, Based on Bruining’s hypothesis^5^ (Fig. 2A), if a time series is generated by a critical-like mechanism:

- An inhibition-dominated dynamics is expected to have 0 *< fE/I <* 1
- An excitation-dominated dynamics will yield 1 *< fE/I <* 2.
- A balanced E/I dynamics will show *fE/I*≈1

For estimating the $w_{amp}(t)$ and $w_{nF}(t)$ function, we used a window size *t* of 50 oscillatory cycles (e.g., 5 seconds at 10 Hz) with 60% overlap between adjacent windows.

##### Estimate bistability

The *BiS* index of a narrow-band power time series (*i.e.*, squared amplitudes) derives from model comparison between a bimodal or unimodal fit of its probability distribution function (PDF). A large *BiS* means that the observed pdf is better described as bimodal, and when *BiS* 🡪 0 the PDF is better described as unimodal. We followed the approach used in ^7,8^ to compute *BiS*. First, to find the PDF of power time series (*R^2^*) of a SEEG contact, the empirical *R^2^* was partitioned into 200 equal-distance bins and the number of observations in each bin was tallied. Next, maximum likelihood estimate was used to fit a single-exponential function (*i.e.*, the square of a Gaussian process follows an exponential PDF):

$P_{x}(x)=\gamma e^{-\gamma x}$	(3)

and a bi-exponential function:

$P_{x}\left( x \right)=\delta\gamma_{1}e^{-\gamma_{1}x}-(1-\delta)\gamma_{2}e^{-\gamma_{2}x}$	(4)

where γ_1_ and γ_2_ are two exponents and *δ* is the weighting factor. Next, Bayesian information criterion (*BIC*) was computed for the single- and bi-exponential fitting:

$BIC = ln(n)k - 2ln(\hat{L})$	(5)

where *n* is the number of samples; $\hat{L}$ is the likelihood function; *k* is the number of parameters: *k* = 1 for single-exponential *BIC_Exp_* and *k* = 3 for bi-exponential model *BIC_biE_*. Thus BIC imposes a penalty to model complexity of the bi-exponential model ^9^ because it has two more degrees of freedom (second exponents and the weight δ) than the single exponential model.

Last, the BiS estimate is computed as the log_10_ transform of difference between the two BIC estimates as

*dBIC* = *BIC_Exp_* - *BIC_biE_*	(6)

*BiS = log_10_(dBIC), if dBIC > 0,*

*BiS = 0, if dBIC ≤ 0*

Thus, a better model yields a small BIC value, and BiS will be large if the bi-exponential model is a more likely model for the observed power time series (S. Fig 3 F, I).

Computational models for local ^10,11^ and large-scale^12,13^ dynamics show that bistability emerges only within a subset of the critical regime when the system is governed by state-dependent positive feedback. In contrast, unimodal oscillations span a broader parameter range in the critical regime, consistent with the classical criticality hypothesis^10,14^ (see Fig. 1, Supplementary Fig. 2 of ^14^). The BiS index uses Bayesian information criterion to estimate the bimodality of the oscillation power distribution and thereby provides an empirical estimate of the strength of a hypothetical positive feedback loop.

#### Large-scale phase synchrony assessments

Pairwise synchrony between two SEEG contacts was estimated using the phase-locking value (*PLV*) for 50 narrow-band frequencies (Morlet wavelets, *m* = 7*.*5)^15^, spanning from 2 to 450 Hz with equal spacing on a *log*_10_ scale. We treat each *PLV* matrix as a graph, wherein contacts are *nodes* and synchronizations are *edges*. The complex-valued phase locking value is defined as:

$cPLV\left( A, B \right)=\frac{1}{T}\sum_{t=1}^{T} \left[ e^{i\left( \theta_{A}\left( t \right)-\theta_{B}\left( t \right) \right)} \right]=\frac{1}{T}\sum_{t=1}^{T} \left[ \frac{S_{A}S_{B}^{*}}{\left| S_{A} \right|\left| S_{B} \right|} \right]$	,	(7)

where *T* denotes the number of samples; *θ_A_* and *θ_B_* are the instantaneous phases of signal *A* and *B*; *S_A_* and *S_A_* are complex-valued narrow-band signals from *A* and *B*, and ^*^ denotes the complex conjugate. We used the [Brain Connectivity Toolbox](https://sites.google.com/site/bctnet/) to characterize *first-order and second-order* synchrony. The *first-order* synchrony was assessed with *effective weight* (*We*) and *eigenvector centrality* (*EVC*). *We* of a node is weighted sum of edges connected to the node. *EVC* is a self-referential centrality so that a node has a high *EVC* estimate if its neighbors have high *EVC*. We assessed *second-order* synchrony derivatives using *clustering coefficient* (*Cc*) and *Local efficiency* (*LE*). *Cc* is the fraction of node’s neighbors that are also neighbors of each other. *LE* is the *efficiency* assessed in the neighborhood of a given node, where *Efficiency* is the average inverse shortest pathlength.

Our previous study^14^ found that while these first- and second-order synchrony features are most effective for cohort-level seizure zone (SZ) classification, criticality features outperform them for within-subject SZ classification. We postulated that, unlike local criticality measures, synchrony features are more susceptible to individual variability in brain network topology and technical factors, which may reduce their specificity at the individual level.

Since first- and second-order nodal strengths are derived from PLV estimates, they may be biased by both physiological and technical factors. The spatial extent, network structure, and synchrony strength of the EpiNet vary substantially across individuals. For example, patients with simple focal seizures and well-defined semiology likely have more localized and consistent EpiNet organization than those with multifocal seizures and complex semiology. Moreover, individual synchrony strength is influenced by a subject's operating point in the critical regime^2^. PLV estimates also decrease with increasing frequency and inter-contact distance^16,17^. These sources of variability, compounded by the inherently sparse spatial sampling of SEEG, likely contribute to the reduced specificity of synchrony features at the individual level.

#### Supplementary Figures

##### S. Fig. 1

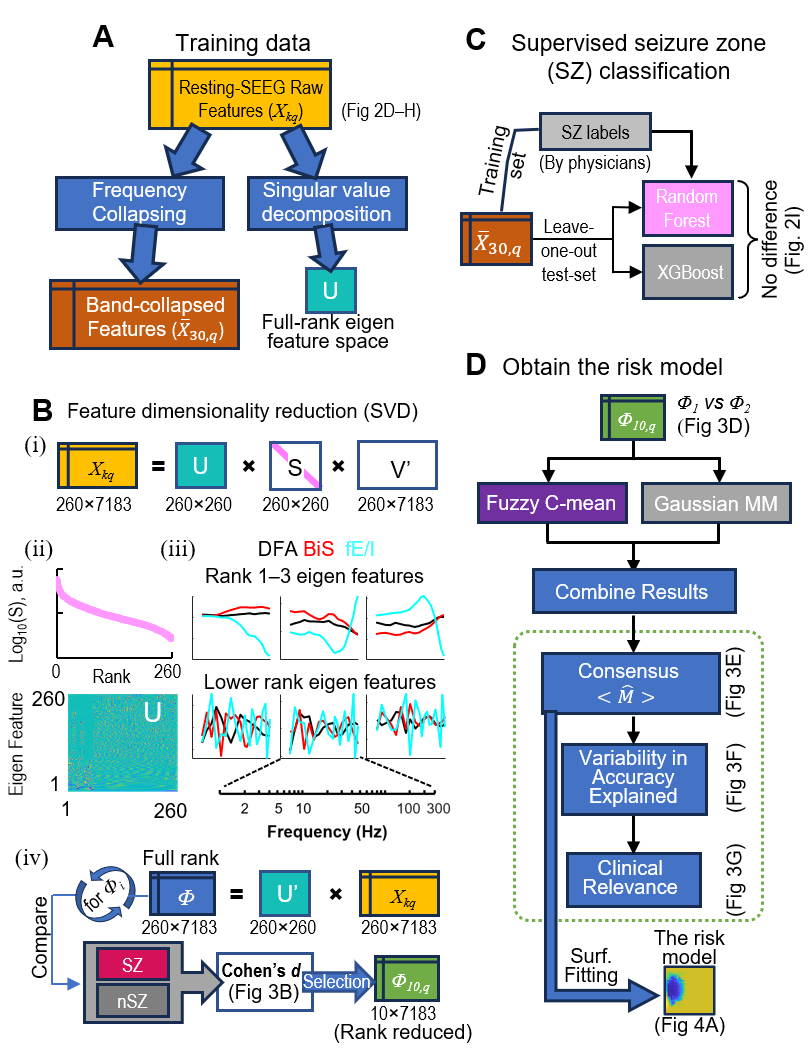

**Additional information about the risk model training.** **(A)** Raw features of the interictal-SEEG from the 64 focal epilepsy patients were collapsed to train supervised classifiers and subject to singular value decomposition for training the seizure-risk model using unsupervised classifiers. **(B)** SVD of the raw features. In practice, the original full matrices of $S\in R^{k\times q}$ and $V^{'}\in R^{q\times q}$ (*k*=260 features; *q*=7,183 contacts) can be ‘thinned’ as shown here. (i) Data dimensionality; (ii) Singular value and the *U* matrix; (iii) Rank 1 to 3 SZ-relevant eigen features; (iv) Eigen-feature selection using clinically identified seizure-zone (SZ). **(C)** Training two supervised classifiers for SZ. **(D)** Pipeline for training the seizure risk model using the rank 1-10 eigen-feature coefficients (*Φ*_10,_ *_q_*).

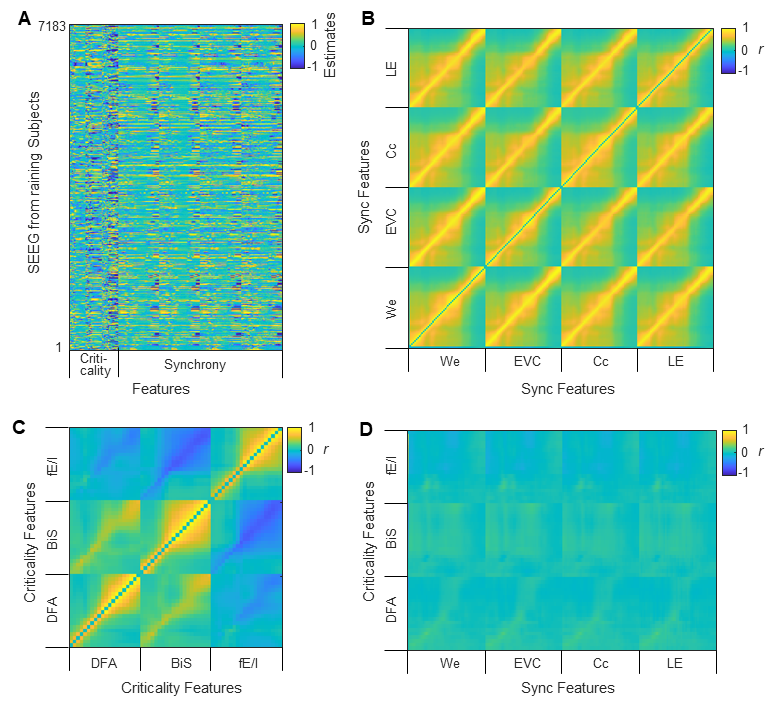

##### S. Fig. 2

**Correlations between narrow-band raw features. (A)** Normalized raw feature values across all contacts (*n* = 7,183) from the training set subjects. **(B–D)** Pairwise Pearson’s correlation coefficients between **(B)** synchrony features (mean *r^2^* = 0.211), **(C)** criticality features (mean *r^2^* = 0.114), and **(D)** synchrony and criticality features (mean *r^2^* = 0.004), and the *r^2^* among synchrony features (B) was significantly greater than that among criticality features (C) (unpaired t-test, -log10(*p*) < 12).

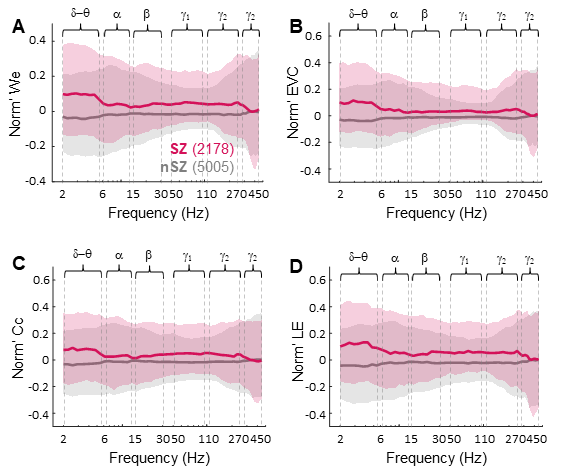

##### S. Fig. 3

**Normalized narrow-band synchrony features for contacts from clinically identified seizure zone (SZ) and non-SZ (nSZ).** **(A)** We: effective weight. **(B)** EVC: eigen vector centrality. **(C)** Cc: clustering coefficient. **(D)** LE: local efficiency. Tick lines: mean; shaded areas: 25% and 75%-tile of all samples (*n* = 7,183).

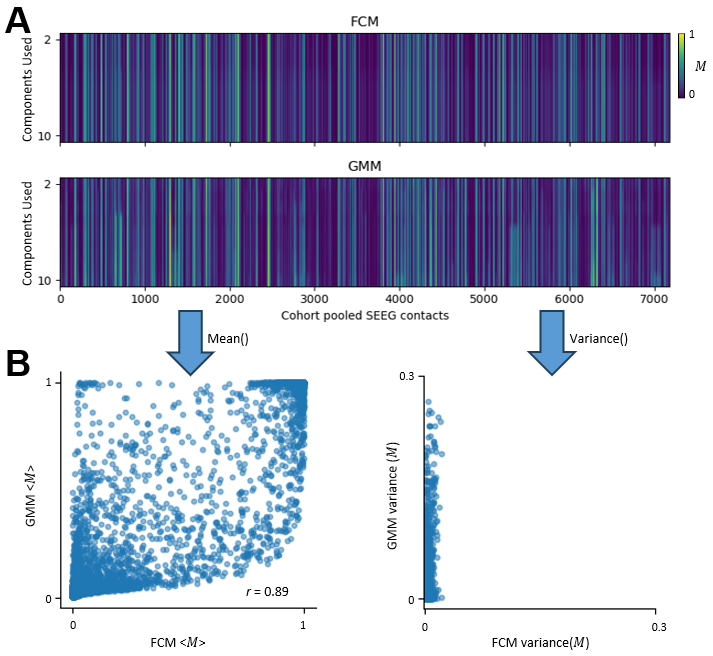

##### S. Fig. 4

**Interim data used to obtain consensus EpiNet-membership for fitting the EpiNet-saliency model. (A)** EpiNet-membership (*M*) assignments for individual SEEG contacts using Fuzzy C-means (FCM, top) and Gaussian Mixture Model (GMM, bottom) across increasing numbers of eigenfeature components (from 2 to 10). **(B)** The mean (left) and variance (right) of membership scores across the tests in (A) are plotted for GMM as a function of those from FCM, where GMM showed large variance in *M* estimates across tests. Markers indicate contacts (*n* = 7,183). The consensus EpiNet-membership is the mean of GMM <*M*> and FCM<*M*> shown in (B) left panel.

### References

1.	Hardstone R, Poil SS, Schiavone G, et al. Detrended fluctuation analysis: A scale-free view on neuronal oscillations. *Front Physiol*. Published online 2012. doi:10.3389/fphys.2012.00450

2.	Fuscà M, Siebenhühner F, Wang SH, et al. Brain criticality predicts individual synchronization levels in humans. *Nat Commun*. Published online 2023:2022.11.24.517800. doi:doi.org/10.1038/s41467-023-40056-9

3.	Palva JM, Zhigalov A, Hirvonen J, Korhonen O, Linkenkaer-Hansen K, Palva S. Neuronal long-range temporal correlations and avalanche dynamics are correlated with behavioral scaling laws. *Proc Natl Acad Sci*. Published online 2013. doi:10.1073/pnas.1216855110

4.	Zhigalov A, Arnulfo G, Nobili L, Palva S, Palva JM. Relationship of Fast- and Slow-Timescale Neuronal Dynamics in Human MEG and SEEG. *J Neurosci*. Published online 2015. doi:10.1523/JNEUROSCI.4880-14.2015

5.	Bruining H, Hardstone R, Juarez-Martinez EL, et al. Measurement of excitation-inhibition ratio in autism spectrum disorder using critical brain dynamics. *Sci Rep*. 2020;10(1). doi:10.1038/s41598-020-65500-4

6.	Diachenkoa M, Sharma A, Smit DJA, et al. Functional excitation- inhibition ratio indicates near- critical oscillations across frequencies. *Imaging Neurosci*. Published online 2024.

7.	Freyer F, Roberts JA, Ritter P, Breakspear M. A Canonical Model of Multistability and Scale-Invariance in Biological Systems. *PLoS Comput Biol*. Published online 2012. doi:10.1371/journal.pcbi.1002634

8.	Roberts JA, Boonstra TW, Breakspear M. The heavy tail of the human brain. *Curr Opin Neurobiol*. Published online 2015. doi:10.1016/j.conb.2014.10.014

9.	Wit E, van den Heuvel E, Romeijn JW. ’All models are wrong. ’: An introduction to model uncertainty. *Stat Neerl*. Published online 2012. doi:10.1111/j.1467-9574.2012.00530.x

10.	Wang SH, Siebenhühner F, Arnulfo G, et al. Critical‐like brain dynamics in a continuum from second-to first‐order phase transition. *J Neurosci*. 2023;43(45):7642-7656. doi:10.1523/JNEUROSCI.1889-22.2023

11.	Cowan JD, Neuman J, van Drongelen W. Wilson–Cowan Equations for Neocortical Dynamics. *J Math Neurosci*. 2016;6(1):1-24. doi:10.1186/s13408-015-0034-5

12.	di Santo S, Villegas P, Burioni R, Muñoz MA. Landau–Ginzburg theory of cortex dynamics: Scale-free avalanches emerge at the edge of synchronization. *Proc Natl Acad Sci*. 2018;115(7):E1356-E1365. doi:10.1073/pnas.1712989115

13.	Dumeur M, Wang SH, Palva JM, Ciuciu P. Multifractality in critical neural field dynamics. Published online 2023:1-6. http://arxiv.org/abs/2312.03219

14.	Wang SH, Arnulfo G, Nobili L, et al. Neuronal Synchrony and Critical Bistability: Mechanistic Biomarkers for Localizing the Epileptogenic Network. *Epilepsia*. Published online 2024. doi:doi.org/10.1101/2023.05.21.541570

15.	Palva JM, Wang SH, Palva S, et al. Ghost interactions in MEG/EEG source space: A note of caution on inter-areal coupling measures. *Neuroimage*. Published online 2018. doi:10.1016/j.neuroimage.2018.02.032

16.	Arnulfo G, Wang SH, Myrov V, et al. Long-range phase synchronization of high-frequency oscillations in human cortex. *Nat Commun*. 2020;11(1). doi:10.1038/s41467-020-18975-8

17.	Dotson NM, Hoffman SJ, Goodell B, Gray CM. A Large-Scale Semi-Chronic Microdrive Recording System for Non-Human Primates. *Neuron*. 2017;96(4):769-782.e2. doi:10.1016/j.neuron.2017.09.050
